# Transcription factor networks disproportionately enrich for heritability of blood cell phenotypes

**DOI:** 10.1101/2024.09.09.611392

**Authors:** Jorge Diego Martin-Rufino, Alexis Caulier, Seayoung Lee, Nicole Castano, Emily King, Samantha Joubran, Marcus Jones, Seth R. Goldman, Uma P. Arora, Lara Wahlster, Eric S. Lander, Vijay G. Sankaran

## Abstract

Most phenotype-associated genetic variants map to non-coding regulatory regions of the human genome. Moreover, variants associated with blood cell phenotypes are enriched in regulatory regions active during hematopoiesis. To systematically explore the nature of these regions, we developed a highly efficient strategy, Perturb-multiome, that makes it possible to simultaneously profile both chromatin accessibility and gene expression in single cells with CRISPR-mediated perturbation of a range of master transcription factors (TFs). This approach allowed us to examine the connection between TFs, accessible regions, and gene expression across the genome throughout hematopoietic differentiation. We discovered that variants within the TF-sensitive accessible chromatin regions, while representing less than 0.3% of the genome, show a ∼100-fold enrichment in heritability across certain blood cell phenotypes; this enrichment is strikingly higher than for other accessible chromatin regions. Our approach facilitates large-scale mechanistic understanding of phenotype-associated genetic variants by connecting key *cis*-regulatory elements and their target genes within gene regulatory networks.

## Introduction

Human genetic studies in population-based cohorts have uncovered tens of thousands of genetic variants associated with complex diseases and phenotypes. With increasingly large sample sizes and diverse ancestry, the fraction of heritability explained is beginning to approach saturation [1–4]. However, genetic mapping alone does not reveal the functional mechanisms by which the variants affect the traits, limiting biological understanding and the development of therapeutic strategies [5].

One approach is to use base editing, in both cell lines and primary cells, to assess the effect of variants on an isogenic background and connect them with target genes and molecular pathways implicated in human disease. While there has been recent progress in achieving high-throughput base editing [6–10], systematic analysis of all variants of interest is not currently feasible.

Another approach is based on the fact that most trait-associated variants are located in non-coding regulatory elements of the genome, where they alter how these elements bind transcription factors (TFs) and related proteins and thereby regulate gene expression [11–13]. A powerful way to study the cellular mechanisms influenced by genetic variation could thus be to modulate sets of TFs and simultaneously read out the genome-wide consequences on both chromatin accessibility of *cis*-regulatory elements and gene expression **(Fig. 1A)**. To assess many TFs and many cell states involved in physiology and disease [14,15], such studies should be performed at single-cell resolution and should be feasible in disease-relevant primary cells [16,17].

**Figure 1.**
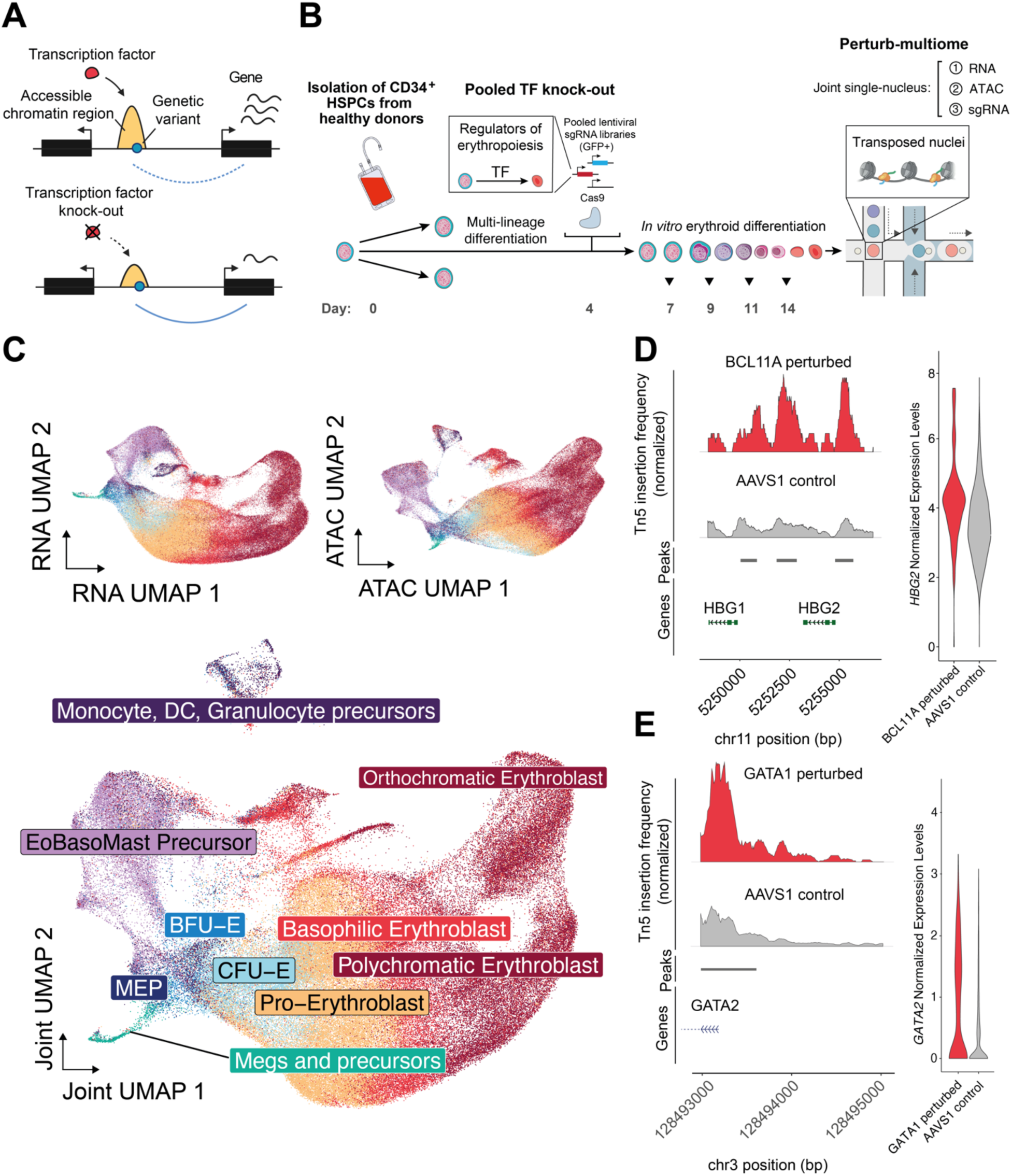
Perturb-multiome: pooled CRISPR screens with coupled single-cell RNA and chromatin accessibility readouts to profile primary human hematopoiesis. **(A)** Top: simplified representation of a basic gene regulatory network in which a transcription factor (TF) modulates chromatin accessibility of a *cis*-regulatory element (enriched in trait- or disease-associated genetic variation) and its linked gene. Bottom: changes in accessibility and expression following experimental TF knock-out. (**B**) Schematic of the experimental setup for pooled TF knock-out with multimodal single-cell readouts. Days spanned since hematopoietic stem and progenitor cells (HSPCs) thawing are noted at the bottom. (**C**) Top left: UMAP reduction using RNA measurements collected from all single cells in the experiment. Top right: UMAP reduction using chromatin accessibility measurements collected from all single cells in the experiment. Bottom: UMAP reduction using a weighted-nearest neighbor graph [18] to integrate RNA and chromatin accessibility peak information from the same single cells. Cell types were annotated using a human bone marrow dataset [19] (**Materials and Methods**). (**D**) Left: normalized Tn5 insertion frequency in BCL11A perturbed cells (perturbation score > 2, Materials and Methods) and control AAVS1 cells. Chromatin accessibility peaks are denoted below and were called using all cells in the experiment. Right: *HBG2* gene expression levels in BCL11A perturbed cells and AAVS1 control cells. (**E**) Left: normalized Tn5 insertion frequency in GATA1 perturbed cells (perturbation score > 2, **Materials and Methods**) and control AAVS1 cells. Chromatin accessibility peaks are denoted below and were called using all cells in the experiment. Right: *GATA2* gene expression levels in GATA1 perturbed cells and AAVS1 control cells.

Here, we report a method, Perturb-multiome, for doing so and apply it to study perturbations of key regulatory TFs in primary human hematopoietic stem and progenitor cells (HSPCs), which can then be differentiated to produce mature red and other blood cells. We find that the regulatory regions whose chromatin accessibility is sensitive to perturbation of these TFs (“TF-sensitive regulatory regions”) comprise less than 0.3% of all regions in the genome, but are enriched by nearly 100-fold for SNPs associated with certain blood-cell phenotypes. Our framework can be applied to human diseases and phenotypes beyond hematopoiesis to advance the understanding of genetic variants and the mechanisms underlying inherited risk for complex diseases.

## Results

### Pooled transcription factor perturbation screens to uncover coordinated chromatin accessibility and RNA alterations

Perturb-multiome applies a library of CRISPR-based perturbations to a cell population and recovers three distinct readouts from each single cell: (1) the identity of the genetic perturbation (sgRNA), (2) the effect of chromatin accessibility at *cis*-regulatory elements (scATAC-seq), and (3) the effect of expression of genes (scRNA-seq) **(Fig. 1A, B)**.

To make Perturb-multiome highly effective on primary cells, we designed the method to overcome challenges in gene-editing efficiency and perturbation identification. To achieve high editing efficiency with minimal toxicity, we deliver pooled sgRNAs via an optimized CROP-seq vector and electroporated recombinant Cas9 ribonucleoprotein [6,20] **(Fig. 1B, Supp. Fig. S1A–C, Materials and Methods)**. To achieve high efficiency in identifying perturbations, we developed a biotin-based enrichment strategy **(Materials and Methods, Supplementary Protocol).**

We applied Perturb-multiome to primary human HSPCs, which can be differentiated to faithfully recapitulate hematopoietic differentiation *in vitro* [6,21,22]. We perturbed 19 TFs, ranging from well-known master hematopoietic regulators to others with suggested roles in hematopoiesis **(Supp. Fig. S3A)** [23]. For each TF, we delivered three different sgRNAs, targeting distinct positions in the coding sequences **(Supp. Table S1 and Materials and Methods)**. We profiled a total of 137,604 cells across four time points. With modest sequencing depth, we were able to (i) detect the identity of the sgRNA in 92.6% of cells; (ii) from the chromatin-accessibility readout, detect an average of 27,774 ATAC fragments and 10,803 unique ATAC peaks per cell; and (iii) from the gene-expression readout, detect an average of 15,970 RNA molecules from 4,129 unique genes per cell **(Supp. Fig. S2A–H)**. Across the cell population, we detected a total of 230,083 unique ATACs peaks and 33,415 expressed transcripts.

We focused on the process of red blood cell production, or erythropoiesis, because nearly half of blood cell trait-associated variants have associations with the erythroid lineage, and genetic variation explains a substantial portion (between 7.19-26.95%) of the heritability of these phenotypes (**Supp. Fig. S5A, Materials and Methods**) [1,2,11]. Following a brief initial stage of multi-lineage differentiation to enable some differentiation to non-erythroid fates, we promoted erythroid differentiation of HSPCs. Cells were assayed at days 7, 9, 11 and 14. We recovered a wide range of cells states, from early common progenitors and non-erythroid lineages (megakaryocyte, monocyte, dendritic cell, and granulocyte precursors) to terminal erythroid precursors that closely mimic cell states present in human bone marrow [19] (**Fig. 1C, Supp. Fig. S1D–J, Materials and Methods)**.

The Perturb-multiome data faithfully identified various well-characterized gene regulatory mechanisms in hematopoiesis. For example, cells with perturbations in the BCL11A TF, a repressor of fetal hemoglobin and a regulator of the fetal-to-adult hemoglobin switch [24,25], displayed both increased chromatin accessibility at the fetal hemoglobin promoters and increased gene expression at the fetal hemoglobin genes (*HBG1/2*), compared to controls **(Fig. 1D)**. Similarly, cells with perturbations in the GATA1 TF showed significantly higher promoter accessibility and gene expression at GATA2 (**Fig. 1E)**, consistent with the negative feedback loop between GATA1 and GATA2 [26,27]. These results illustrate the ability of Perturb-multiome to obtain fine-grained insights into physiologically relevant processes directly in primary human cells.

### Identification of perturbed cells and transcription factor-sensitive accessible chromatin regions and genes

As a first step in our analysis, we sought to characterize (1) the gene-editing efficiency of each sgRNA and (2) the perturbation level actually achieved in each individual cell, which can vary among cells receiving the same sgRNA.

To characterize gene-editing efficiency, we used pooled single-cell genotyping to assess the editing efficiency for sgRNAs used in the screen **(Supp. Fig. S3B, Materials and Methods)** [6]. We relied on genotyping rather than the mRNA level of the targeted gene, because some CRISPR-Cas9 edits that disrupt protein-coding regions do not affect mRNA levels. We analyzed 17,665 single cells in droplets, performing 55 multiplexed PCRs in each — corresponding to the human genomic sites targeted by sgRNAs, as well as 10 regions within the integrated lentiviral genome to identify the sgRNA in the cell — to estimate the editing efficiency (**Fig. 2A, left, Supp. Fig. S3C,D)**.

**Figure 2.**
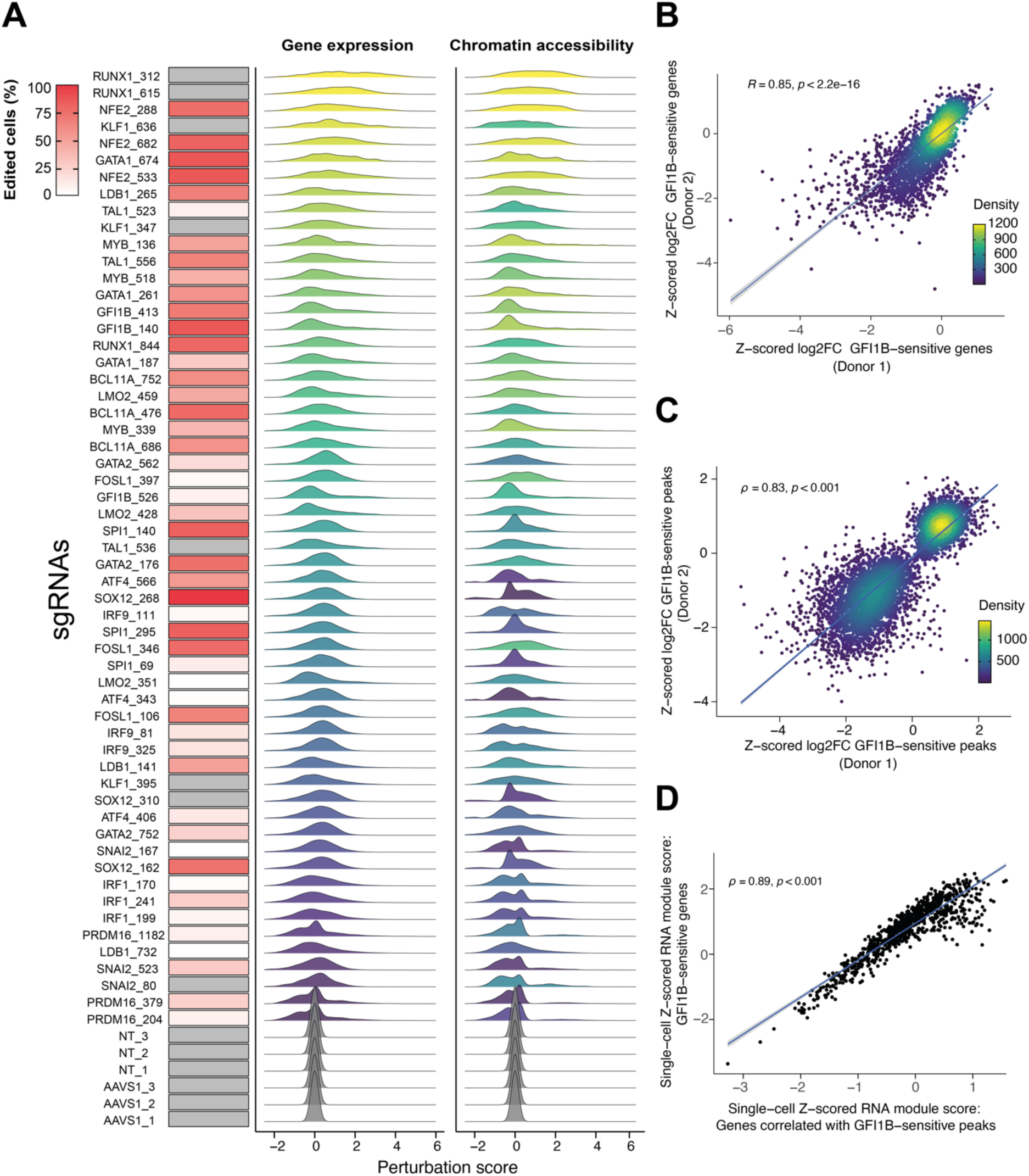
Widespread gene expression and chromatin accessibility changes in response to transcription factor perturbations. **(A)** Left column: percent of edited cells from pooled single-cell genotyping experiments for each sgRNA **(Supp. Fig. S3, B to D, Materials and Methods)**. Amplicons with missing data (not covered by single-cell multiplexed PCR) and non-targeting (NT) guides are shown in grey. Middle column: per-cell perturbation score distribution computed on single-cell RNA counts. Right column: per-guide perturbation score distribution computed on single-cell chromatin accessibility peak counts. Each sgRNA is ordered along the color gradient based on the median perturbation score across all cells with that guide, for the RNA or chromatin accessibility modalities. **(B)** Scatter plot of the Z-scored log_2_FC for RNA gene expression levels (shown are statistically significant genes after multiple hypothesis testing correction) between cells with one of the GFI1B-targeting sgRNAs and AAVS1-targeting control sgRNAs from two independent pooled TF Perturb-multiome screens in two different HSPC donors. The Spearman correlation coefficient ρ is shown, as well as a regression line with confidence intervals. **(C)** Scatter plot of the Z-scored log_2_FC for ATAC peak counts (shown are statistically significant genes after multiple hypothesis testing correcting) between cells with one of the GFI1B-targeting sgRNAs and AAVS1-targeting control sgRNAs for two independent pooled TF Perturb-multiome screens in two different HSPC donors. The Spearman correlation coefficient <u>ρ</u> is shown, as well as a regression line with confidence intervals. **(D)** Scatter plot of the single-cell Z-scored RNA module score computed using GFI1B-sensitive genes (defined as in panel B) and the single-cell Z-score RNA module score computing using genes correlated with GFI1B-sensitive ACRs (defined as in panel C, **Materials and Methods**), for cells with a Z-scored perturbation score greater than 1. The Spearman correlation coefficient ρ is shown, as well as a regression line with confidence intervals.

To characterize the perturbation level achieved by an sgRNA in each individual cell, we defined “perturbation scores” for both the ATAC and RNA data by comparing the chromatin accessibility and gene expression profiles of the cell to those from nearest-neighbor control cells, which represent a similar differentiation state **(Fig. 2A, middle and right, Supp. Fig. S3F, Materials and Methods)** [28]. The perturbation scores were well correlated with sgRNA editing efficiency **(Supp. Fig. S3E)**. The exceptions included two TFs (GATA2 and SPI1) for which we achieved strong knock-down (confirmed by protein levels), but saw low perturbation scores in erythroid lineages; these TFs have their strongest effects in non-erythroid lineages **(Supp. Fig. S3G)** [29,30].

We then defined TF-sensitive accessible chromatin regions (ACRs) and TF-sensitive genes by using a linear model that incorporated the perturbation level achieved in each single cell **(Materials and Methods)**. We found that roughly 21.8% of the ACRs (50,114/230,083) **(Supp. Table S2, Supp. Fig. S5B)** and 26.0% of expressed genes (8,694/33,415) (**Supp. Table S3**) were affected by at least one TF perturbation. Of these TF-sensitive results, 32% of accessible chromatin regions and 22% of genes responded to only a single TF perturbation **(Supp.** Fig. S5, G, I). To assess robustness of the data, we examined an independent biological replicate (using HSPCs from a different donor and sgRNAs targeting different locations in the coding sequences of each TF) and found the results were highly correlated, both qualitatively and quantitatively, with average correlation of >0.9 for TFs perturbed in both replicates (**Fig. 2B,C**, **Supp. Fig. S4A)**.

### Properties of transcription factor-sensitive elements

Having used Perturb-multiome to separately analyze changes in chromatin accessibility and gene expression, we next examined the correlation between these features in single cells carrying a specific sgRNA vs. control guides to define ACR-gene linkages **(Fig. 3A, Materials and Methods)**. To assess the ACR-gene linkages, for each cell, we defined two scores: (i) the first equal to the average expression of the TF-sensitive genes for a given TF and (ii) the second equal to the average expression of the genes correlated with TF-sensitive ACRs. We found strong concordance between these scores (**Fig. 2D, Materials and Methods**).

**Figure 3.**
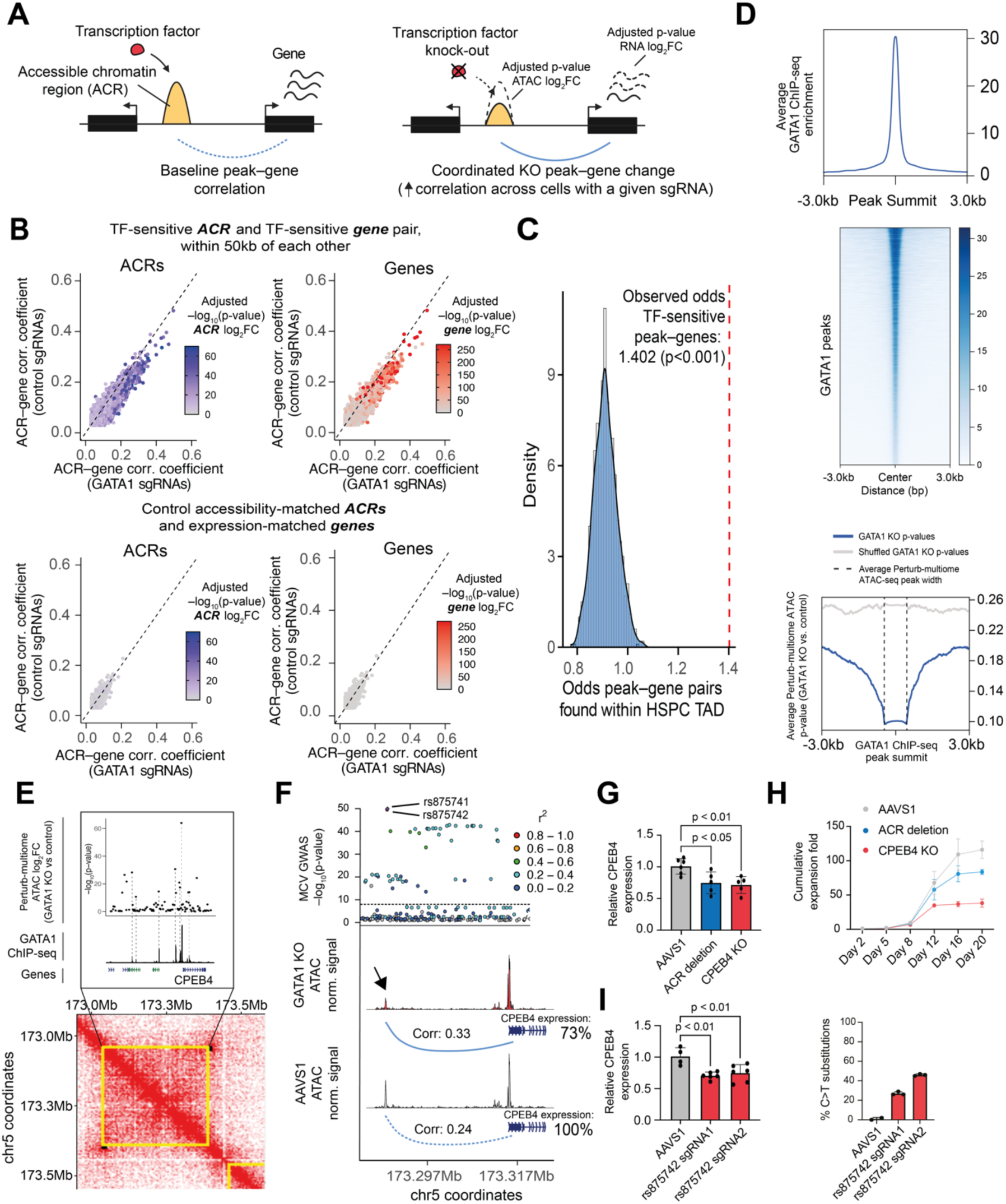
Genome-wide characterization of TF-sensitive elements. **(A)** Simplified representation of the metrics analyzed in this figure for a basic TF-ACR-target gene regulatory network. **(B)** Scatter plots of ACR–gene correlations across cells transduced with control sgRNAs and cells transduced with GATA1-targeting sgRNAs. The diagonal represents the correlation values for ACR–gene pairs with equal correlation coefficients among cells transduced with control sgRNAs and cells transduced with GATA1-targeting sgRNAs. Left panels: the color scale represents the p-value adjusted for multiple hypothesis testing of the ATAC ACR log2FC between GATA1-targeting and control sgRNAs. Right panels: the color scale represents the p-value adjusted for multiple hypothesis testing of the gene expression log2FC between GATA1-targeting and control sgRNAs. Top panels: ACR–gene correlations for pairs comprising a TF-sensitive ACR and a TF-sensitive gene within 50kb from one another. Bottom panels: ACR–gene correlations for pairs comprising a non-TF-sensitive ACR and a non-TF-sensitive gene (each selected among a distribution of non-sensitive elements with similar accessibility levels to their TF-sensitive counterparts) within 50kb of each other. **(C)** Distribution of the odds that control ACR-gene pairs (each non-TF-sensitive, each selected among a distribution of non-sensitive elements with similar accessibility levels to their TF-sensitive counterparts) fall within an HSPC TAD[31], built from random sampling from this distribution 1000 times **(Materials and Methods).** The red dotted line represents the observed odds for TF-sensitive ACR-gene pairs. The p-value represents how many of the sampled control observations are larger than the observed odds. **(D)** Top: profile plot of independent GATA1 ChIP-seq enrichment surrounding called ChIP-seq peaks from day 12 erythroid progenitors[32]. Middle: heatmap plot for GATA1 ChIP-seq reads surrounding GATA1 peaks. Bottom: profile plot of Perturb-multiome adjusted -log10(pvalues) for the log2FC of accessibility for ACRs in cells with GATA1-targeting sgRNAs compared to control centered on GATA1 ChIP-seq peaks as defined in the top panel. **(E)** Analysis of Perturb-multiome differential accessibility results within an HSPC TAD. Shown are the Perturb-multiome adjusted - log10(pvalues) for the log2FC of accessibility for ACRs in cells with GATA1-targeting sgRNAs compared to control, GATA1 ChIP-seq signal [32], gene annotations highlighting the GATA1-sensitive CPEB4 gene, and chromosome loops from Hi-C data [31]. TADs are shown in yellow and loop anchors in black **(Materials and Methods)**. **(F)** Top: locus zoom plot showcasing -log10(pvalues) for variants associated with mean corpuscular volume (MCV) [2]. The color scale represents the squared correlation (r^2^) between rs875741 and other neighboring alleles. Middle and bottom: normalized ATAC signal in cells with GATA1-targeting and AAVS1-targeting cells, respectively. The TF-sensitive ACR with the most significant change compared to control is highlighted with an arrow. The Spearman’s rank correlation coefficient (ρ) between this ACR and CPEB4 is shown, as well as the mean CPEB4 expression relative to control. **(G)** Bar plots of the relative CPEB4 expression for HSPCs edited with a control dual cut in the AAVS1 locus, a deletion of the putative *CPEB4* enhancer on chromosome 5, or a CPEB4 knock-out, and subsequently differentiated into the erythroid lineage. Data for day 14 of erythroid differentiation, for two distinct HSPC donors. **(H)** Cumulative expansion fold for HSPCs edited with a control dual cut in the AAVS1 locus, a deletion of the putative *CPEB4* enhancer on chromosome 5, or a CPEB4 knock-out, and subsequently differentiated into the erythroid lineage. **(I)** Left, bar plots of the relative CPEB4 expression for HSPCs edited with the TadCBE cytosine base editor precomplexed with AAVS1-targeting control sgRNAs or two different sgRNAs targeting rs875742. Data for day 14 of erythroid differentiation, for two distinct HSPC donors. Right, gene editing efficiencies for one of the donors shown in the left panel.

First, we examined the effect of TF perturbation on the correlation between pairs of ACRs and genes lying within 50 kb of one another. For the most TF-sensitive ACRs and TF-sensitive genes, the correlation (at the single-cell level) between the pairs showed a significant increase; this result was seen at all 6 erythroid TFs for which there was a substantial number of such pairs **(Supp. Fig. S4B)**. For example, the most significant GATA1-sensitive ACRs and genes had higher correlations in cells expressing a GATA1-targeting sgRNA compared to controls **(Fig. 3B, top).** In contrast, control non-TF-sensitive gene–ACR pairs did not display this behavior (**Fig. 3B, bottom, Materials and Methods**). This is consistent with coordinated regulatory changes at the ACRs and genes, and it suggests the possibility of using this method to reconstruct gene regulatory networks on a genome-wide scale.

Second, we considered ACR-gene pairs within 2 Mb that showed correlated changes and examined their three-dimensional proximity, based on Hi-C data from human HSPCs **(Materials and Methods)** [31]. We found that significant TF-sensitive ACR–gene pairs had higher odds of being found within topologically-associated domains (TADs) in HSPCs (OR 1.54, p < 0.001) (**Fig. 3C).**

Third, we observed that TF-sensitive genes exhibited lower RNA polymerase II pausing compared with non-TF-sensitive genes with similar expression levels **(Supp. Fig. S4C, Materials and Methods)** [33].

Fourth, we were interested in whether TF-sensitive ACRs occurred near binding sites for the perturbed TF. To this end, we analyzed a published dataset of ChIP-seq data for two of our TFs, GATA1 and NFE2, in primary human proerythroblasts **(Materials and Methods)** [32]. Importantly, we found that TF-sensitive ACRs in the regions surrounding GATA1 and NFE2 ChIP-seq peaks showed stronger significance levels (that is, higher – log_10_(p-values)) **(Fig. 3D, Supp. Fig. S4D)**.

We then focused on one of the ACR-gene pairs with the highest increase in correlation upon GATA1 perturbation. This pair, located at 5q35.2, involves the *CPEB4* gene and an ACR 25 kb upstream of the gene’s TSS and lies within a TAD in HSPCs. *CPEB4* plays a role in mouse erythropoiesis, but has not been studied in humans [34]. The ACR has two variants in strong linkage disequilibrium with each other that are associated with multiple blood cell traits [1,2]. Strikingly, the most significant TF-sensitive ACRs within the TAD were strongly correlated with GATA1 ChIP-seq peaks **(Fig. 3E)** [2]. Besides having one of highest changes in gene-ACR correlation, independently, *CPEB4* and the ACR each also had the most significant changes in response to GATA1 perturbation within the TAD **(Fig. 3F).** These observations suggest that GATA1 binds at the accessible chromatin region to modulate expression of *CPEB4*.

To test this functional hypothesis, we used CRISPR-Cas9 to delete the ACR in primary cells. The deletion resulted in a decrease of *CPEB4* expression during erythroid differentiation **(Fig. 3G, Supp. Fig. S4E).** Furthermore, perturbation of *CPEB4* and, to a lesser extent, deletion of the regulatory element resulted in impaired terminal erythroid differentiation, consistent with the variant associations within the ACR with erythroid mean corpuscular volume, mean corpuscular hemoglobin, red blood cell distribution width, and mean corpuscular hemoglobin concentration, and prior studies on the role of *CPEB4* in mouse erythropoiesis **(Fig. 3H)** [34]. We next examined the effect of editing one of the variants, using the recently developed base editor TadCBE [35] (**Materials and Methods).** Strikingly, this single-nucleotide change reduced the expression of *CPEB4* in primary erythroblasts **(Fig. 3I**). Together, these results illustrate how coupling readouts of TF perturbation, gene expression, and chromatin accessibility in the same single cells can identify TF-sensitive ACR-gene pairs with important roles in human erythropoiesis and can help systematically pinpoint associations between phenotypes and variants discovered by GWAS.

### Transcription factor-sensitive *cis*-regulatory elements are massively enriched in the heritability of blood cell traits

Encouraged by the CPEB4 example, we sought to examine more broadly how TF-sensitive regulatory regions were related to naturally-occurring genetic variation underlying blood-cell phenotypes and diseases. We began by noting that TF-sensitive ACRs are under greater constraint (lower rate of genetic variation) [36] compared to the non-accessible genome **(Fig. 4A, Materials and Methods)**. Furthermore, for several TFs, the average constraint for TF-sensitive ACRs was higher than for non-TF-sensitive ACRs. This suggested that the TF-sensitive ACRs may play key, conserved functional roles in hematopoiesis.

**Figure 4.**
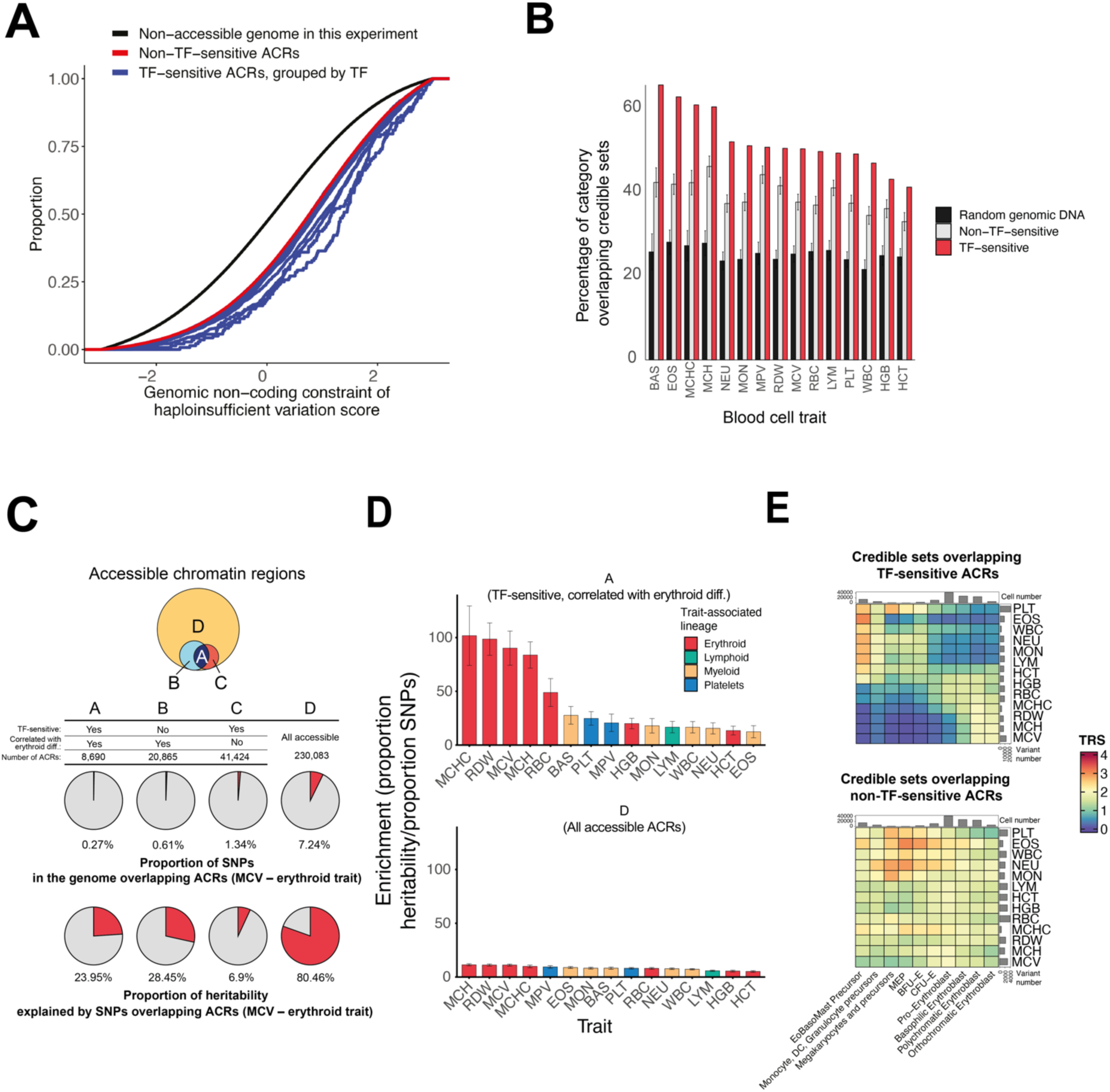
Characterization of genetic variation within TF-sensitive *cis*-regulatory elements. **(A)** Cumulative distribution of the genomic non-coding constraint of haploinsufficient variation score [36], colored by functional category. **(B)** Barplots for the percentage of 95% credible sets overlapping TF-sensitive ACRs, for each blood cell trait. Error bars represent the standard deviation of 100 sampling events of non-TF-sensitive accessible ACRs (“non-TF-sensitive”) or of any genomic region (“random genomic DNA”) **(Materials and Methods). (C)** Top: area proportional Venn Diagrams representing the number of total ACRs. Middle: proportion of SNPs in the genome that overlap ACRs for mean corpuscular volume (MCV). Bottom: proportion of heritability explained by SNPs overlapping ACRs. **(D)** Barplots of the enrichment (proportion of heritability divided by the proportion of SNPs) for two of the four functional categories defined in panel C (categories A and D). Traits are colored by the respective associated lineage. **(E)** Heatmaps of the Trait Relevant Score (TRS) **(Materials and Methods)** for variants belonging to credible sets overlapping TF-sensitive ACRs (top) and not overlapping TF-sensitive ACRs (bottom). Similar enrichments were observed using only variants from the 95% credible sets directly overlapping with the aforementioned elements (data not shown).

We explored whether genetic variants associated with blood-cell phenotypes were enriched in the TF-sensitive ACRs, by examining how credible sets for each trait overlapped with TF-sensitive ACRs, non-TF-sensitive ACRs, and random genomic regions. We found that, averaged across traits, the proportion of regions that contained a variant in a credible set for a trait was much higher for TF-sensitive ACRs (50.59%) than for non-TF-sensitive ACRs (39.25%) and random genomic regions (22%) (**Fig. 4B).** When we examined the results for individual TFs and traits, we found striking differences. The highest proportion was 65.75%, for TF-sensitive ACRs containing variants in credible sets for basophil number **(Fig. 4B, Supp. Fig. S5F, H, Supp. Fig. S6, Materials and Methods)**.

For each of the blood cell traits, we also examined the proportion of heritability explained by SNPs in various regions, using partitioned linkage disequilibrium score regression (LDSC) **(Materials and Methods).** We examined three types of ACRs: (i) TF-sensitive ACRs active during erythroid differentiation, (ii) non-TF-sensitive ACRs active during erythroid differentiation, and (iii) all ACRs observed observed in both erythroid and non-erythroid cells, **(Materials and Methods, Supp. Fig. S5C)**. Strikingly, while TF-sensitive ACRs active during erythroid differentiation comprise <0.3% of the genome, heritability in these regions showed ∼100-fold enrichment for several traits — that is, ∼25% of the heritability lies in this small proportion of the genome **(Fig. 4C, D)**. In contrast, the non-TF-sensitive ACRs active during erythroid differentiation showed lower enrichment (up to ∼40-fold enrichment for certain traits) **(Supp. Fig. S5D)**. The set of all ACRs comprises much more of the genome (7.24%) and explains more of the heritability (∼79%), but the enrichment is much lower (∼11-fold) **(Fig. 4C, D)**. Notably, the TF-sensitive ACRs were also significantly enriched in TF binding motifs compared to all ACRs active solely during erythroid differentiation **(Supp. Fig. S5E).** Furthermore, we observed that credible sets of GWAS variants that lie within the TF-sensitive regions showed clear patterns of cell-type specific enrichments for their respective traits, unlike other variants **(Fig. 4E, Materials and Methods)**. These results suggest a critical role for TF-sensitive ACRs and the associated variants lying within them in explaining the heritability of blood cell phenotypes.

Collectively, we observe that chromatin regions with accessibility changes in response to transcription factor perturbation are disproportionately enriched for phenotype-relevant genetic variation. More broadly, if this is true for other complex diseases and phenotypes, it suggests that systematic mapping of TF-mediated gene regulatory networks will accelerate in variant-to-function mapping.

## Discussion

Understanding the molecular and functional consequences of non-coding variation is essential to dissect the mechanisms underlying complex diseases and phenotypes. These insights have proven key for the development of new diagnostic and therapeutic interventions, as recently achieved in hemoglobin disorders by emerging gene therapies [12,22,32,34,36]. Scalable approaches to functionalize regulatory elements and link them to large sets of candidate trait-associated genetic variation are critical for the progress towards these goals, especially in human tissues with complex mixtures of cells.

We performed pooled perturbations of hematopoietic TFs in primary human hematopoietic cells to obtain massively parallel single-cell readouts of their effects in hundreds of thousands of genomic regions and transcripts. Analysis of genome-wide TF-sensitive regions led to the identification of a tiny subset of the genome (<0.3%) that is highly enriched in heritability for blood cell phenotypes, relative to hematopoietic- or erythroid-specific accessible chromatin. Notably, these regions overlapped an average of 51% of credible sets for blood cell traits, and had up to a 100-fold enrichment in heritability for certain cell-traits. In addition, variants overlapping these regions displayed specific genetic enrichments, consistent with their functional importance in specific hematopoietic populations.

Several of the TFs studied are master regulators of hematopoietic differentiation and their perturbation results in developmental blocks [37–40]. We nonetheless obtained fine-grained observations of the effects of transcription factors at each stage by using analytical approaches that weigh comparisons by the perturbation scores of each single cell with respect to nearest-neighbor control cells, thereby leveraging the asynchronous nature of hematopoietic differentiation and overcoming the variable efficiencies and effects of each perturbation.

The TF perturbation approach is modular and compatible with alternative strategies to modulate gene expression [41–43]. Endogenous tagging of transcription factors with degrons or related approaches could be used to explore the immediate consequences of TF degradation [44–47]. However, this strategy is not currently applicable to large numbers of regulators and in HSPCs isolated from human donors [48]. While we focused on the effects of a single perturbation per cell, future efforts using multiplexed perturbations of regulators could be used to discern the combinatorial logic of transcription factor regulatory control [49,50].

The methods we employ to dissect the properties of TF-sensitive networks (i.e., ChIP-seq, PRO-seq, Hi-C) are independent from the methods used to define TF-sensitive elements (i.e., transcription factor perturbation). A fraction of the gene-ACR linkages changed concomitantly in their expression, chromatin accessibility, and correlation in response to perturbation. These TF-sensitive networks are likely to harbor additional genes and associated regulatory elements with still-to-characterized critical hematopoietic functions.

We focused our analysis on the study of TF-sensitive regions and likely causal variants, which allowed us to study a larger fraction of phenotype-relevant variation beyond those that directly disrupt TF-binding motifs [11]. With continued progress in genome editing and cost-efficiency in single-cell analysis, these regions will be excellent candidates for saturation mutagenesis to identify functional base pairs and pinpoint causal variants.

More broadly, we envision that the identification of TF-sensitive gene regulatory networks through perturbation approaches in complex primary cell contexts will help in prioritizing and understanding the mechanisms underlying a wide range of human diseases and phenotypes across many tissues and organs.

## Supporting information

Supplementary Protocol

## Acknowledgments

We are grateful to the members of the Sankaran and Lander Laboratories, as well as Dr. Amy Guillaumet and Dr. Amrita Sule for valuable discussions regarding this work. J.D.M.R. was supported by fellowships from La Caixa Foundation (ID 100010434), the Rafael del Pino Foundation, and the American Society of Hematology. This work was supported by the Broad Institute Gene Regulation Observatory (V.G.S.), the New York Stem Cell Foundation (V.G.S.), a gift from the Lodish Family to Boston Children’s Hospital (V.G.S.), the Howard Hughes Medical Institute (V.G.S.) and National Institutes of Health (NIH) grants R01 DK103794, R01 CA265726, R01CA292941, and R01 HL146500 (V.G.S.).

## Author contributions

Conceptualization, J.D.M.R., A.C., and V.G.S; Formal analysis, J.D.M.R., A.C., E.K., S.R.G., E.S.L., and V.G.S.; Funding acquisition, J.D.M.R, and V.G.S; Investigation, J.D.M.R., A.C., S.L., N.C., E.K., S.J., S.R.G., U.A., L.W., E.S.L., and V.G.S.; Methodology, J.D.M.R., A.C., and V.G.S; Software, J.D.M.R., A.C., and E.K.; Resources, S.R.G., E.S.L., and V.G.S.; Supervision, V.G.S.; Visualization, J.D.M.R., and A.C.; Writing – original draft, J.D.M.R., A.C., and V.G.S; Writing – review & editing, J.D.M.R., A.C., E.S.L., and V.G.S. with input from all authors.

## Data and code availability

Raw and processed data have been deposited at GEO and will be available at the time of publication. The corresponding code is available in our github repository (https://github.com/jmartinrufino/perturb_multiome/)

## Materials and Methods

### Lentiviral vector

We employed a modified CROP-seq vector for efficient single-cell perturbational screens in hematopoiesis that we previously described [6]. The vector has been deposited in Addgene (https://www.addgene.org/203321/).

### Lentiviral production

Lentiviral particles were produced as previously described[6]. Briefly, Human Embryonic Kidney-293T cell line (ATCC, CRL-3216), was cultured in DMEM Life Technologies, 1965-118) supplemented with 10% FBS (BioTechne, S11550) and 1% penicillin-streptomycin (Life Technologies, 15140-122). When they reached ∼70% confluency, cells were co-transfected with the pooled CRISPR library along with VSV-G and pDelta 8.9 packaging vectors overnight, using FuGene lipofection (Promega, E2691). Media was changed the following morning to DMEM supplemented with 20%FBS, and viral supernatant was first collected 24 hours later and kept at 4°C. Media was subsequently replaced and harvested again 48 hours later. After cold centrifugation at 1,500 rpm and filtration through 0.45µm PVDF membrane (Millipore, SCHVU01RE) to remove cell debris, supernatants were pooled and ultracentrifuged at 24,000 rpm, 4°C for 1:30h (Beckman Coulter SW32Ti). The pellet was ultimately resuspended with StemSpanTM SFEM-II media (Stemcell Technologies, #09655) and stored at -80°C until further usage.

### Erythroid culture

5 million CD34^+^ cells were thawed in 1X PBS (Gibco, 10010-023) supplemented with 1% FBS, centrifuged at room temperature, x350g for 5min, and resuspended in 10mL of hematopoietic stem cell (HSC) expansion media consisting of StemSpanTM SFEM-II media (Stemcell technologies, #09655) supplemented with 1% L-Glutamine (Thermo Fisher Scientific, 25-030-081), 1% penicillin/streptomycin (Life Technologies, 15140-122), 50ng/mL human TPO (PeproTech, 300-18) and 100x CC100 cytokines (StemCell Technologies, 02690). Erythroid differentiation was then induced in 3 steps as previously described [6]. On day 3, cells were washed in 50mL 1X PBS and resuspended in phase I erythroid media at a density of 10^5^ cells/mL. Phase I media is IMDM based (Life Technologies, 12440-061) and supplemented with 3% AB human serum (Atlanta Biologicals, S40110), 2% AB human plasma (SeraCare, 1810-0001), 1% penicillin/streptomycin (Life Technologies, 15140-122), 10µg/mL recombinant insulin (Lilly, NDC 0002-8215-01), and 3 IU/mL heparin (Hospira, NDC 00409-2720-01), 200µg/mL human holo-transferrin (Sigma-Aldrich, T0665), 1ng/mL recombinant human IL-3 (Peprotech, 200-03), 10ng/mL human recombinant Stem Cell Factor (SCF, Peprotech, 300-07), 3IU/mL recombinant human erythropoietin (EPO, epoetin alpha, Amgen, NDC 55513-267-10). Cells were spinfected with the CRISPR library on day 2 overnight, then washed and resuspended in the same phase I media. Cells were nucleofected with Cas9 on day 4 and resuspended in phase I media until day 7. On day 7, recombinant human IL-3 was withdrawn from the media and cells were cultured until day 11 in phase II media. On day 11, holo-transferrin concentration was increased to 1mg/mL and both IL-3 and SCF were removed from the media, and cells were cultured in phase III media until the end of the culture. Media was freshly renewed every 2-3 days, and cells were thoroughly washed in 1X PBS (Gibco, 10010-023) to remove the cytokines at each step of the culture.

### Lentiviral transduction

On day 2, CD34+ cells were infected at a density of 10^6^ cells per mL in HSC expansion media supplemented with 8µg/mL polybrene (Sigma Aldrich, TR-1003-G). Freshly thawed lentiviral particles were added at a MOI of 0.3-0.5, and cells were spinfected at 2,000 rpm for 1:30h at 37°C. Cells were kept overnight at 37°C. The next morning, infected cells were washed in 50 times their volume of 1X PBS and resuspended in Phase-I erythroid media.

### Cas9 electroporation

Cells were nucleofected on day 4, 2 days after spinfection, after verifying the expression of VEX reporter by flow cytometry. To nucleofect 15 million cells, 10µl of 62µM Alt-R S.p. HiFi Cas9 Nuclease V3 (IDT, 1081061) was diluted in 21µl of 1X PBS and then combined with a non-targeting sgRNA in a swap strategy that has proven to increase the editing efficiency in HSCs for cutting Cas9 [20], for 20min at room temperature. In the meantime, cells were washed twice with 1X PBS to remove RNAses from the media, then resuspended in 190µl P3 solution with supplement (Lonza, VPA-1003) and 10µl of 100µM Alt-R Electroporation enhancer (IDT, 1075916). Cell suspension was then dispensed in two 100µl electroporation cuvettes and electroporated in a 4D-Nucleofector X Unit (Lonza) using the DZ-100 program. Immediately after, 500µl of phase-I erythroid media was added to each cuvette and incubated at 37°C for 5min. Then, cells were resuspended in phase-I media at a density of ∼500,000 cells per mL.

### FACS enrichment prior to single-cell genomics

To enrich cells expressing a single gRNA at each time point (days 7, 9, 11 and 14), we sorted cells for expression of VEX (Violet light excited GFP reporter present in the lentiviral vector, **Supp. Fig. S1C**). 10mL of cell suspension was washed in 40mL of 1X PBS (x300g, 5min, room temperature) and the pellet was resuspended in 2mL 1X PBS + 0.04%BSA. 6-8×10^5^ cells were sorted in a 5mL low binding tube (Eppendorf, 0030122356) pre-coated with 1mL of 1X PBS + 0.04%BSA (Sigma-Aldrich, A9418), using a Sony MA900 sorter. Sorted cells were then pelleted at 300g for 5min, at 4°C, resuspended in 250µl of 1X PBS + 0.04%BSA and placed on ice.

### Multiparametric flow-cytometry (FCM) of cell differentiation

At each time point, 10^5^ unsorted cells were washed in 1mL of 1X PBS and resuspended in 100µl of 1X PBS + 0.04%BSA. 5×10^4^ cells were used for unstained and each single cell color control. Cells were stained with a cocktail of antibodies targeting CD123, CD36, CD34, CD71 and CD325a, and 7-AAD or with one of those as a single-color control (**Supp. Table S4**). On day 14, only antibodies targeting CD71, CD235a and 7-AAD staining were used. After 30min of incubation at room temperature in the dark, cells were washed twice with 1mL 1X PBS + 0.04% BSA, and resuspended in 100µl of the same solution. Acquisition was performed on the same Sony MA900 as sorted cells. For intracellular staining, 10^5^ cells were stained with anti-human CD71 and CD235a antibodies, and fixed in 1X PBS + 4% paraformaldehyde (PFA) for 15min at room temperature. After 2 washes in 1X PBS, cells were resuspended in 1X PBS + 2% FBS. To permeabilize, 100% Methanol was added to a final concentration of 90% while gently vortexing and incubated on ice for 10min. After 2 washes, cells were incubated for 30min in the dark at room temperature in 1X PBS + 2% FBS with intracellular antibodies targeting PU.1, GATA-1, or GATA-2 at manufacturer’s recommended dilution. Cells were washed twice and fluorescence was read on a CytoFLEX S cytometer (BD).

### Morphological analysis of erythroid differentiation

At each time point (days 7, 9, 11 and 14), ∼10^5^ sorted cells were washed and resuspended in 150µl of 1X PBS + 50% FBS and centrifuged in a Cytospin 4 centrifuge (Thermo Fisher Scientific) at 500rpm for 5min, with low acceleration. Slides were air dried, and first stained with May-Grünwald solution (Sigma Aldrich, MG1L) for 5 min, rinsed 4 times in MilliQ water for 30s per wash, then stained with 1:20 dilution of Giemsa solution (Sigma Aldrich, 32884) for 15min and rinsed 6 times in MilliQ water for 30sec per wash. Slides were air dried overnight and imaged on a Mica instrument (Leica microsystems).

### Precision nuclear run-on sequencing (PRO-seq)

CD34+ cells from 2 different XY donors were cultured in erythroid phase I media for 8 days, and electroporated with 1nmol GATA1 or AAVS1 control sgRNAs pre-complexed with 1nmol Cas9 15min at RT. 5-10 million cells were electroporated in P3 solution with P3 Primary Cell 4D-Nucleofector™ LV Kit L, program DZ-100 (V4LP-3002), and resuspended in phase II erythroid media. Editing efficiency was 80% as assessed by Sanger sequencing, and flow cytometry showed a 3-fold decrease in intracellular GATA1-PE mean fluorescence intensity in the CD71^+^/CD235a^-^ progenitor population. Cells were then harvested 24h post-electroporation and processed following the Harvard Nascent Transcriptomics Core cell permeabilization protocol for PRO-seq [51]. Briefly, cells were harvested on ice in 5mL low binding tubes (Eppendorf, 0030108310), and washed twice in cold PBS at 400g, 4°C, for 5min. After removing cold PBS, cells were mixed in 200µl of Wash Buffer (W) to resuspend a single cell suspension. Cells were permeabilized in 4mL of Permeabilization Buffer (P) for 5min on ice. Cells were then centrifuged at 400g, 4°C, for 5min, to remove the supernatant and immediately resuspended in 1mL of W-buffer to dilute any remaining P-buffer. After gently mixing, 4mL of W-buffer was added and cells were washed at 400g, 4°C, for 8min. The supernatant was removed and cells were resuspended in 2×100µl F-buffer in a 1.5mL low binding tube (Eppendorf, 022431021). Cells were counted and Freezing Buffer (F) was adjusted to a cell concentration of 5-10×10^6^/mL, flash frozen on dry ice. PRO-seq was then performed as previously published [52].

### Assessment of enhancer cutting efficiency

Genomic DNA was extracted from ∼100,000 cells using QuickExtract™ DNA Extraction Solution (Biosearch Technologies, QE0905T). The chromosome 5 enhancer locus was PCR amplified using Platinum™ II Hot-Start PCR Master Mix (2X) (ThermoFisher scientific, 14000013) for 30 cycles, and its deletion was verified on E-Gel™ 2% Agarose gel with SYBR™ Safe DNA stain (ThermoFisher scientific, A42135). Relative quantification of fragments was assessed with the Agilent High Sensitivity DNA Kit (5067-4626) on a Agilent 2100 Bioanalyzer instrument. The 900bp/1,650bp molarity ratio was estimated using the bioanalyzeR R package. Deletion efficiency was confirmed with Sanger sequencing on the column-purified PCR fragments (QIAquick PCR Purification Kit, Qiagen, 28104), using the forward PCR primer for sequencing. Primers are referenced in **Supp. Table S5**.

### Real-Time quantitative Polymerase Chain Reaction (RT-qPCR)

Total RNA was extracted from around 100,000 cells using column purification with removal of genomic DNA using DNAse I (RNEasy Micro Kit, Qiagen, 74004). After Reverse Transcription (iScrpit^TM^, BioRad, 1708841), 10ng of complementary DNA (cDNA) was used for SYBR green quantitative PCR (LightCycler® 480 SYBR Green I Master, 04887352001) and detected on CFX384 Touch Real-Time PCR Detection System, BioRad). Primers are listed in **Supp. Table S5**.

### Cytosine base editing

The TadCBE cytosine base editor was purified using the procedure we previously reported [6]. We designed a codon optimized version of TadCBE with an N-terminal His tag [35]. The plasmid for purification has been deposited in Addgene (https://www.addgene.org/225093/).

We performed two sequential electroporations, on day 3 and day 4 after thawing, using the DZ-100 electroporation program in a 4D-Nucleofector X Unit (Lonza), as previously described for HSPC electroporation with base editors [6]. Immediately after, 100µl of phase I erythroid media was added to each cuvette and incubated at 37°C for 5min. Then, cells were distributed in phase I media at a density of ∼500,000 cells per mL. Cuvettes were again washed with 75µl of phase I erythroid media to retrieve any cells from the cuvette.

### Perturb-multiome

At each time point, 160,000 sorted cells were processed in 4 technical replicates. Nuclei were extracted following 10× Genomics recommendations for nuclei isolation for single cell multiome CG000365 RevB, with the recommended buffers. All centrifugation steps were carried in a swinging bucket centrifuge to reduce cell loss. 40,000 sorted cells were placed in a low binding tube (Eppendorf), volume was adjusted to 50µl with 1X PBS + 0.04% BSA, and cells were centrifuged at 300 rcf for 5min at 4°C. 45µl of supernatant was removed and replaced with 45µl of pre-chilled Lysis buffer containing 10mM Tris-HCl pH 7.4, 10 mM NaCl, 3 mM MgCl2, 1% BSA, 0.1% Tween 20, 0.1% NP-40, 0.01% digitonin, 1 mM DTT, 1 U/mL Protector RNase inhibitor, and gently mixed 3 times before incubating 3 min on ice. Then, 50µl of chilled wash buffer containing 10 mM Tris-HCl pH 7.4, 10 mM NaCl, 3 mM MgCl2, 1% BSA, 0.1% Tween 20, 1 mM DTT, 1 U/mL Protector RNase inhibitor, was added to each tube without mixing, and tubes were centrifuged at 500 rcf for 5 min at 4°C. 95µl of supernatant was then removed and replaced with 45µl of pre-chilled diluted nuclei buffer containing 1X final concentration of 20X Nuclei Buffer, 1 mM DTT, 1 U/mL Protector RNase inhibitor. Nuclei were centrifuged at 500 rcf for 5 min at 4°C, and supernatant was carefully removed in totality. Nuclei pellet was resuspended in 7µl of the same chilled nuclei buffer, and 1µl of the nuclei suspension was stained with 1:10 dilution of 1 mg/mL Propidium Iodide (Life Technology, P3566) and manually counted on a hemocytometer (INCYTO, DHC-N01-2). Nuclei quality was assessed visually on a Nikon Eclipse TS100 microscope after diluting 1µl of the nuclei suspension in 4µl of chilled nuclei buffer and stained with 5µl of 0.4% Trypan blue solution (Invitrogen, T10282). Immediately after counting, 16,100 nuclei per replicate were processed into multiome protocol to target 10,000 nuclei recovery. Transposition, GEM generation and barcoding, and libraries construction were prepared following 10× Genomics protocol for Chromium Next GEM Single Cell Multiome ATAC + Gene Expression CG000338 Rev E. The four replicates were processed in parallel at each time point, in the same Next GEM Chip J (10× Genomics, PN-1000230). The GEM emulsion was then stored at -80°C for a few days until all time points were harvested, and the 16 samples were then processed together for clean-up and further steps. The validation experiment was processed exactly in the same way, but only 2 replicates per time point were processed in parallel.

### Pooled single-cell genotyping

To determine the editing efficiency at single cell level, we designed primer pairs to amplify ∼300bp genomic regions around the predicted cutting site of each sgRNA, as well as 9 primers targeting regions of the integrated CROP-seq vector **(Supp. Fig. S3B, Supp. Table S5)**. Two days after thawing, 2 million CD34+ cells were spinfected with the pooled-CRISPR library at a MOI ∼0.3-0.5 following the method described in the corresponding section of the manuscript and incubated overnight at 37°C. Cells were washed the next morning then resuspended in phase-I of erythroid media. On day 5, 8 million cells were electroporated with Cas9 ribonucleoprotein complexed with a non-targeting sgRNA, following the protocol further detailed. Cells were then resuspended in phase I of erythroid media until day 8. On day 8, 900,000 Vex+ cells were FACS sorted following the protocol detailed in the corresponding section of the manuscript and concentrated to 3,000 cells /µl in 1X PBS + 0.04% BSA and placed on ice. 2 replicates of 100,000 cells were immediately processed using the Tapestri single-cell DNA sequencing V3 (Mission Bio) following manufacturer’s recommendations (User Guide QRC v3_MB05-0010_rev2).

### CROP-seq transcript enrichment

A step by step protocol has been included in **Supplementary Protocol.** To improve the detection of sgRNAs at single cell level, we amplified CROP-seq transcripts containing the sgRNA information using a 2-steps polymerase chain reaction (PCR), from the cDNA amplified in 10× multiome workflow. First, 10ng of cDNA from each sample was amplified with NEBNext® Q5U® Master Mix (NEB, M0597S), with a biotinylated forward primer (/5Biosg/UAUAGTGACTGGAGTTCAGACGTGTGCTCTTCCGATCTCGATTTCTTGGCTTTATATATCT TGTG) and a modified phosphorothioate reverse primer to protect from nuclease cleavage (CTACACGACGCTCTTCCGAT*C*T), with the following parameters: 98°C for 30s, then 8 cycles at 98C for 15s for denaturation, 69°C for 15s for primer annealing, 72°C for 20s for fragment elongation, and a terminal elongation step at 72°C for 120s. PCR1 fragments were cleaned up with 1X SPRIselect beads (Beckman Coulter B23318) and eluted in Buffer EB (Qiagen, 19086). Then, biotinylated PCR1 fragments were enriched by Dynabeads™ MyOne™ Streptavidin C1 (Thermo Fisher scientific, 65001) following manufacturer’s recommendations. Briefly, after 4 washes in WB buffer, PCR fragments were incubated with 0.5X pre-washed Dynabeads™ for 15min at room temperature, on a rotating mixer (HulaMixer, Thermo Fisher scientific), washed 3 times in WB buffer, and incubated in a master mix of 15uL Tris-EDTA buffer solution pH 7.4 (Sigma Millipore, 93302) and 0.5uL of Thermolabile USER (Uracil-Specific Excision Reagent) II Enzyme (NEB, M5508S) for 30min at 37°C on a rotating mixer. PCR1 fragments were then cleant-up with 1X SPRIselect beads and eluted in buffer EB for the second PCR. PCR2 primers included universal Illumina sequencing adaptors P5 and P7 with a different index for each sample, and amplification was carried out with Q5® High-Fidelity DNA polymerase (NEB, M0492L) as following: initial denaturation at 98°C for 30s, followed by 26-30 cycles (determined by qPCR) at 98°C for 15s for denaturation, 69°C for 15s for primer annealing, 72°C for 20s for elongation, and a terminal elongation step at 72°C for 120s. PCR2 fragments were finally cleaned-up with 1X SPRIselect beads and resuspended in buffer EB for sequencing. Primer sequences are references in **Supp. Table S5**.

### Sequencing

Quantification and quality of all libraries were assessed on a Bioanalyzer 2100 using High-Sensitivity DNA Reagent Kit (Agilent, 5067-4626). Single-cell gene expression and ATAC replicates were separately pooled to 2nM, and final concentration was assessed by qPCR using KAPA-quantification kit (Roche, KK2602). Both libraries were sequenced using a Novaseq 6000 S4 kit (Illumina) in pair-ended mode. The CROP-seq transcript enrichment library was sequenced with a NextSeq 1000/2000 P3 100 cycles (Illumina) in pair-ended mode, with a custom primer for read 2 (CGATTTCTTGGCTTTATATATCTTGTGGAAAGGACGAAACACCG). The single cell genotyping library was sequenced with a NextSeq 1000/2000 P2 300 cycles (Illumina) in pair-ended mode.

### Perturb-multiome data processing

Custom scripts for the analyses described in the subsequent sections can be found in this paper’s Github respository.

Raw bcl files were demultiplexed using bclconvert v4.0.3. Gene expression and ATAC libraries were processed using Cellranger ARC 2.0.2. CROP-seq guide enrichment libraries were processed jointly with gene expression libraries using Cellranger v7.0.1. Matrices from all modalities were merged downstream in R. Using Seurat v4, standard processing of scRNA-seq data was performed with *FindVariableFeatures, ScaleData, RunPCA, FindNeighbors, FindClusters* and *RunUMAP* [18]. For the ATAC data, standard processing using Signac and *RunTFIDF, FindTopFeatures, RunSVD* and *RunUMAP* was performed [53]. To create the weighted nearest neighbors graph, *FindMultiModalNeighbors* was used and then UMAP was run on the joint space. Cell types were annotated using human bone marrow data [19].

### Pooled single cell data processing

Sequenced fastq.gz files were demultiplexed using bclconvert v.4.0.3 and processed using the Tapestri Pipeline v3, which comprises adapter trimming, alignment, cell barcode error correction, cell identification among droplets and variant calling, followed by MissionBio’s Genome Editing Solution.

Mutational frequencies in **Fig. S3D** are a weighted frequency of the times that a given type of mutation was seen across reads for one cell barcode, which was then averaged across cells.

### Perturbation score calculation

An initial set of differentially expressed genes was identified by comparing cells with a given guide to their nearest neighbor non-targeting cells (in a similar differentiation state) using Mixscale [28]. This defines a Z-scored perturbation score for each single-cell. That score was then used as a weight in a linear regression model to predict target gene expression levels, such that cells with higher perturbation scores are given a higher weight. In the end, if a perturbed gene controls a target gene, the term associated with that score in the model will be significantly different from 0. To avoid circularity in this strategy, a leave-one-out approach was used, in which the gene predicted in the regression is taken out from the initial perturbation score computation.

We extended this approach to the ATAC dimension by modeling data as fragment counts, rather than reads [54], and using the LSI reduction to calculate perturbation signatures.

TF-sensitive elements were selected for each transcription factor as those whose p-values were significant following Bonferroni correction for multiple hypothesis testing.

### PRO-seq analysis

Sequencing reads were trimmed to 41bp, and only those with an average quality >20 were retained. Adapters were removed using cutadapt 1.14 and 3’ bases with low quality using –match-read-wildcards -m 20 -q 10. An initial alignment step of R1 reads was performed to the control Drosophila genome spike-in using Bowtie 1.2.2 (with the parameters -v 2 -p 6 --best --un). Unmapped reads were then aligned to the hg38 genome (-v 2 --best), sorted and converted to bedGraph format.

RNA polymerase pausing analysis was performed after counting read counts surrounding active dominant transcription start sites (TSSs) using proTSS (https://github.com/NascentTranscriptionCore/proTSScall) compared to gene bodies. Dominant TSSs and dominant transcription end sites (TESs) were obtained from this 5’ PRO-seq data and our previously published human erythropoiesis RNA-seq dataset [6].

### Gene–ACR correlation analysis

Gene expression and chromatin accessibility correlations were computed across single cells for each sgRNA using *FigR*[55]. Briefly, *FigR* calculates the gene-ACR Spearman correlation across single cells and obtains significance estimates by permuting the correlation coefficients of a set of GC-matched background ACRs. We computed correlations over 50kb or 2Mb for every gene-ACR pair, and retained correlations with a Spearman correlation > 0.03.

To obtain control accessibility-matched ACRs and expression-matched genes, we generated a distribution using non-TF sensitive elements with similar mean and skew to the distribution of TF-sensitive ACRs and randomly sampled an equal number of observations.

### Analysis of the overlap between TF-sensitive genes and genes correlated with TF-sensitive ACRs

We calculated the average expression levels of genes of interest, subtracted by the aggregated expression of control genes with similar expression, using the *AddModuleScore* function from Seurat [18]. We did this for TF-sensitives and for genes correlated with TF-sensitive ACRs. Genes correlated with TF-sensitive ACRs were defined as described in the section “Gene–ACR correlation analysis” above, using gene–ACR pairs with a pvalZ < 0.05 [55].

### Topologically-associated domain (TAD) analyses

We used previously published data of HSPC TADs [31]. We reprocessed data using Juicer Tools 1.19.02 HiCCUPS using default settings to obtain loops and used *arrowhead* to obtain TADs. To obtain control accessibility-matched ACRs and expression-matched genes, we generated a distribution using non-TF sensitive elements with similar mean and skew to the distribution of TF-sensitive ACRs and randomly sampled an equal number of observations. By randomly sampling this distribution 1,000 times, we built a baseline expectation of the probability of a given TF-ACR pair to be found within a TAD.

### GATA1 and NFE2 ChIP-seq analyses

We used previously published GATA1 ChIP-seq data from day 12 primary human erythroblasts[32], and previously published NFE2 ChIP-seq data from GYPA^+^ primary human erythroblasts [56]. We reprocessed the data and mapped reads using BWA 0.7.15. We only kept uniquely mapped reads to the human genome using samtools view -q 1. We then called GATA1 and NFE2 peaks using MACS2. To perform metaplots, we used deeptools *computeMatrix* and *plotProfile.* To compute the profiles of *GATA1* and *NFE2* chromatin accessible TF-sensitive peaks, *computeMatrix* and *plotProfile* were run surrounding GATA1 or NFE2 peaks computed using MACS2.

### Genomic non-coding constraint of haploinsufficient variation (Gnocchi) scores

We obtained genome-wide constraint z-scores from gnomAD 3.1 for 1kb windows and overlapped them with our functional elements (i.e., non-accessible chromatin, TF-sensitive accessible ACRs, and non-TF-sensitive ACRs) using the *GenomicRanges* function.

### Blood cell trait credible set and TF-sensitive element overlap analysis

We computed the number of 95% credible sets overlapping with TF-sensitive ACRs for each blood cell trait from one of the largest GWAS for blood cell traits performed to date, Chen et al. [2]. To build control distributions, we obtained 100 random samples of the same number of non–TF–sensitive ACRs, and 100 random samples of the same number of genomic regions, with the same width as the average width of TF-sensitive ACRs.

### Heritability and LD score regression analyses

We used data from European Ancestry from one of the largest GWAS for blood cell traits performed to date, Chen et al. [2]. We ran stratified LD score regression (LDSC)[57] on our annotations alongside the baseline model for each of the fine-mapped summary statistics to compute heritability estimates and enrichments for each trait (proportion of heritability / proportion of SNPs in functional annotation). We used summary statistics and LD Scores from 1000 Genomes and European ancestry.

To define ACRs correlated with erythroid differentiation, we used Palantir with default settings on the ATAC dimension to identify sets of ACRs that monotonically increased over the course of differentiation (**Supp. Fig. S5C)**[58].

### Motif-enrichment analysis

Analysis of overrepresented motifs in TF-sensitive ACRs correlated with erythroid differentiation compared to ACRs correlated with erythroid differentiation alone was performed using Signac [53]. Briefly, motifs from the JASPAR 2020 database [59] were retrieved and added to the single cell object using the *getMatrixSet* and *AddMotifs* functions. An hypergeometric test was used (*FindMotifs* function) to test the probability of observing a motif at a given frequency by chance in TF-sensitive ACRs correlated with erythroid differentiation, compared to ACRs correlated with erythroid differentiation alone.

### Trait-enrichment score (TRS) calculation

We classified credible sets with a 95% probability of containing the causal GWAS variant in European ancestry as overlapping either TF-sensitive or non-TF-sensitive regions (i.e., the fraction of all accessible ACRs in this experiment that were not TF-sensitive). We then used Scavenge to obtain trait-enrichment scores using variants included within credible sets overlapping TF-sensitive and non-TF-sensitive regions, as previously described[60]. Similar enrichments were observed using only variants from the 95% credible sets directly overlapping with the aforementioned elements.

## Supplementary Figures

**Supplementary Figure 1.**
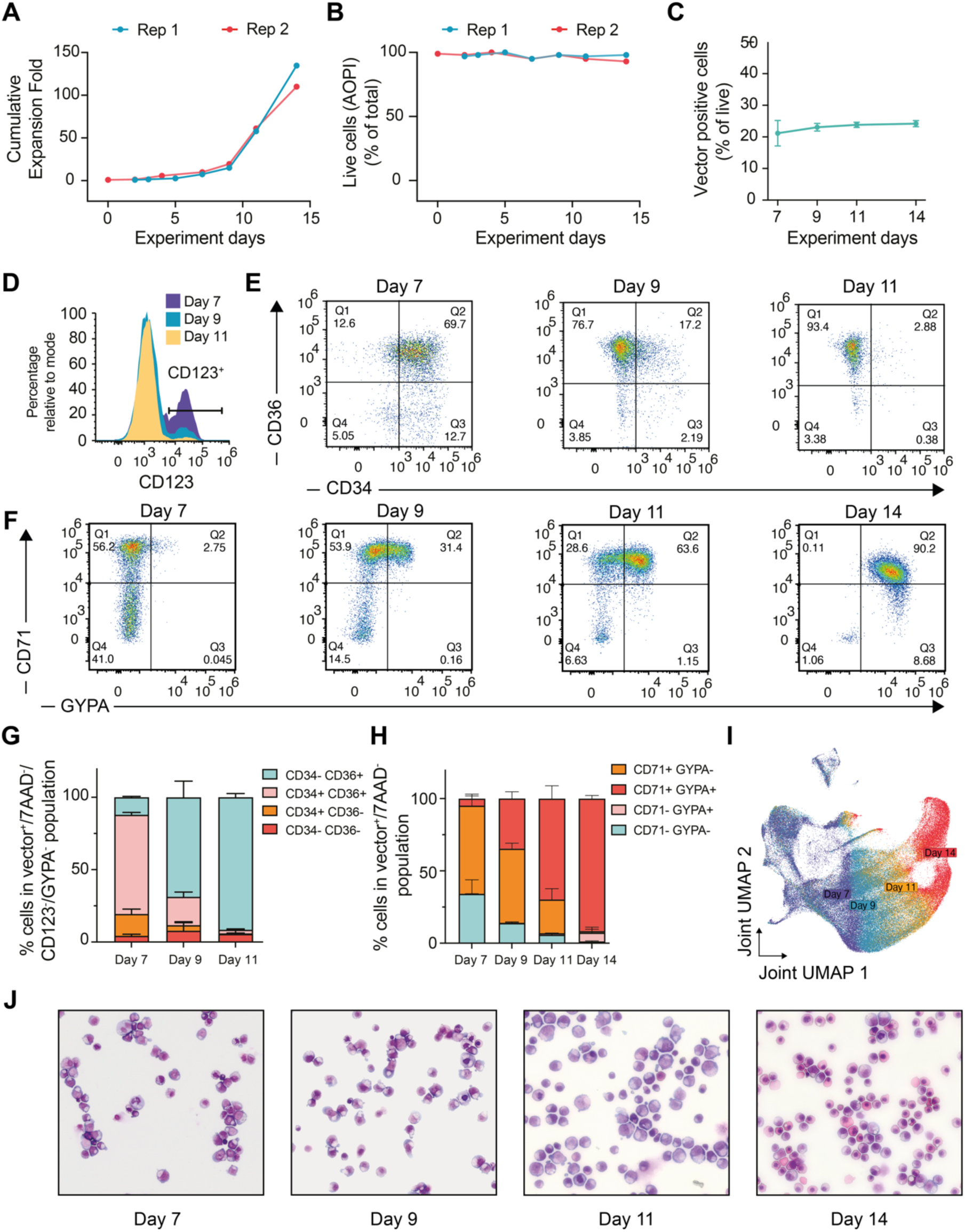
Characterization of erythroid differentiation in cells subjected to Perturb-multiome. **(A)** Cell growth curves of two experimental replicates over the course of the experiment described in Fig. 1. **(B)** Cell viability curves of two experimental replicates over the course of the experiment described in Fig. 1. **(C)** Percentage of differentiating hematopoietic cells expressing the lentiviral vector over each of the experimental time points, following transduction at low multiplicity of infection. Cells were subsequently enriched using FACS to obtain a pure population (>99%) of transduced cells prior to single-cell profiling. **(D)** Histograms of CD123 (IL3RA) protein expression over the course of experimental time points, showing the progressive enrichment of erythroid cells. **(E)** Scatter plots of CD34 and CD36 protein expression over the course of experimental time points, showing the progression through erythroid progenitor stages. **(F)** Scatter plots of GYPA and CD71 protein expression over the course of experimental time points, showing the progressive maturation of erythroid precursors. **(G)** Bar plots for two biological replicates of coordinated changes in CD34 and CD36 protein expression across time points. Mean and standard deviation are displayed. **(H)** Bar plots for two biological replicates of coordinated changes in GYPA and CD71 protein expression across time points. Mean and standard deviation are displayed. **(I)** UMAP reduction using a weighted-nearest neighbor graph [18] to integrate RNA and chromatin accessibility peak information from the same single cells. Cells are colored by the experimental timepoint of profiling. **(J)** Representative microscopy images of sorted cells at each of the experimental time point, used profiled with Perturb-multiome. Cells progress from immature stages with large nucleus and apparent nucleolus (Day 7) to early erythroblasts with large basophilic cytoplasm (Day 9), then decreasing in size and compacting chromatin (Day 11), and ultimately cytoplasm turning polychromatophilic and then acidophilic before enucleation (Day 14) at terminal stages.

**Supplementary Figure 2.**
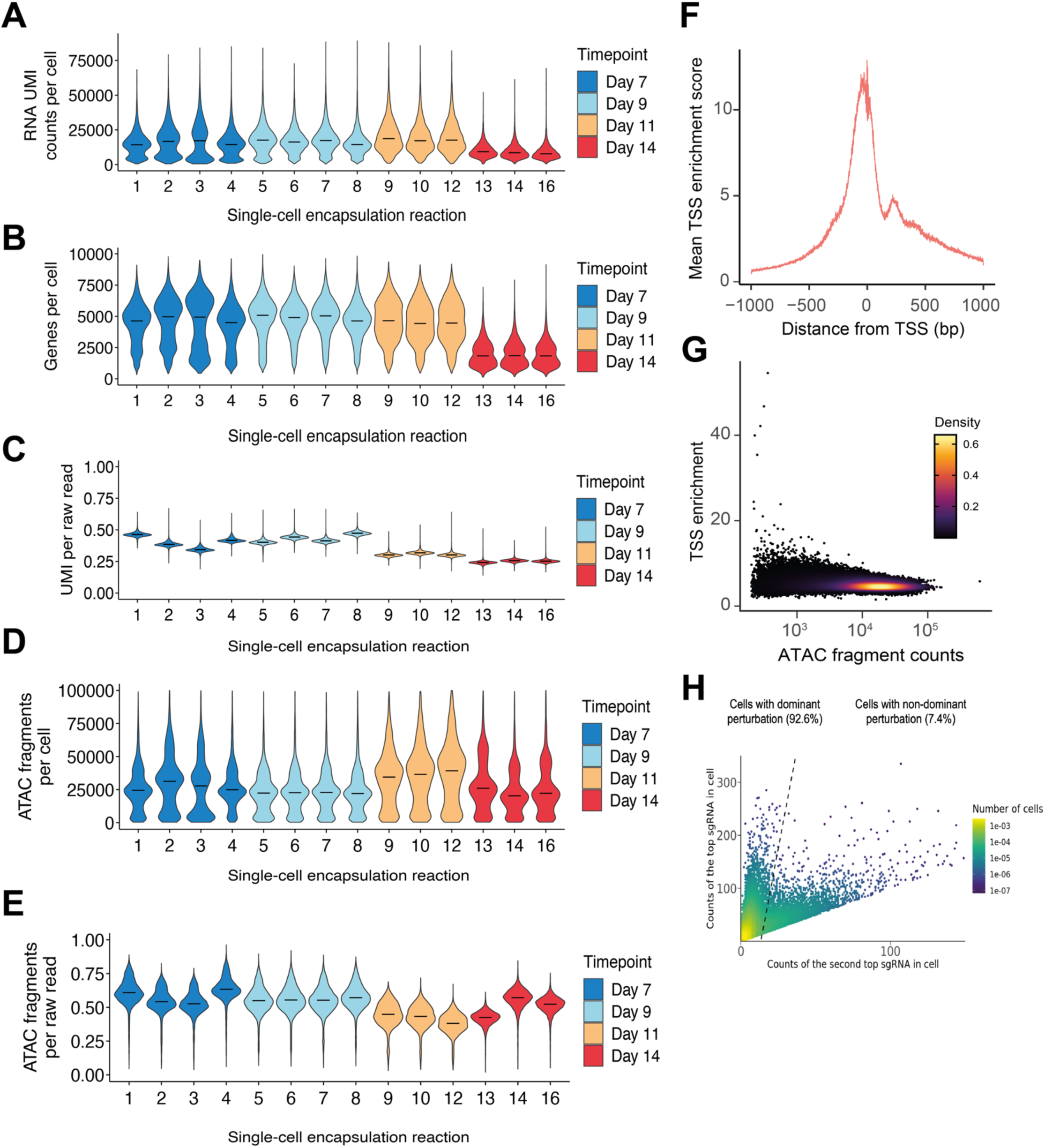
Quality metrics of Perturb-multiome. **(A)** Violin plots of the RNA UMI counts per single cell, grouped by single-cell encapsulation reaction and colored by experimental time point. Horizontal black lines represent the median for each group. **(B)** Violin plots of the genes detected per single cell, grouped by single-cell encapsulation reaction and colored by experimental time point. Horizontal black lines represent the median for each group. **(C)** Violin plots of the ratio between the number of UMI per cell and the number of gene expression raw sequencing reads per cell, grouped by single-cell encapsulation reaction and colored by the time point of experimental collection. Horizontal black lines represent the median for each group. **(D)** Violin plots of the ATAC fragments per single cell, grouped by single-cell encapsulation reaction and colored by experimental time point. Horizontal black lines represent the median for each group. **(E)** Violin plots of the ratio between the number of ATAC fragments per cell and the number of ATAC raw sequencing reads per cell, grouped by single-cell encapsulation reaction and colored by the time point of experimental collection. Horizontal black lines represent the median for each group. **(F)** Mean transcription start site (TSS) enrichment score as a function of the distance to the TSS for all single cells in the experiment. **(G)** Density plot of the per-cell TSS enrichment score as a function of the number of ATAC fragment counts in each cell. **(H)** Density plot of the CROP-seq UMI counts of the top sgRNA in each cell as a function of the CROP-seq UMI counts of the second sgRNA with most counts in each cell.

**Supplementary Figure 3.**
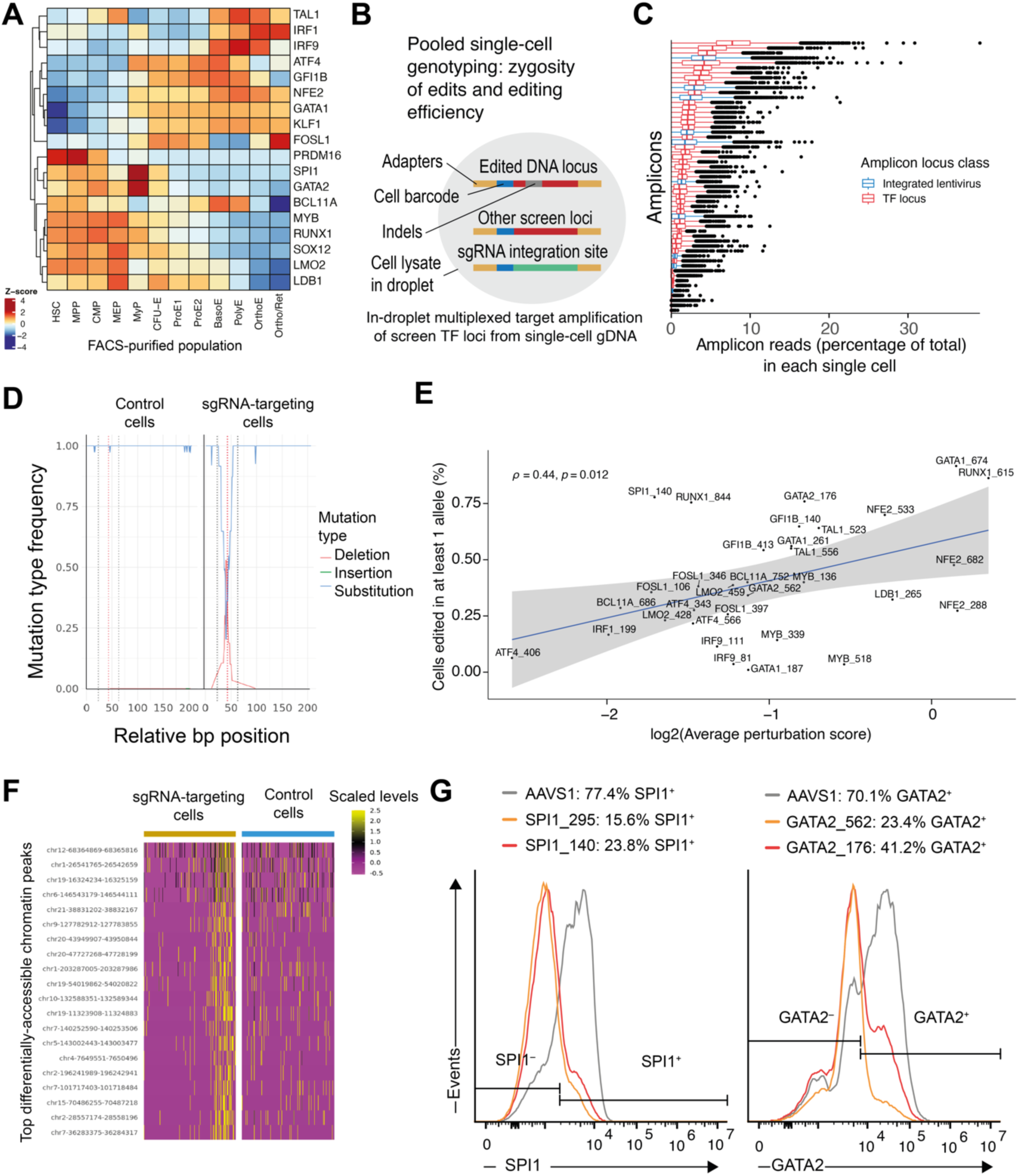
Assessment of transcription factor perturbation efficiencies. **(A)** Heatmap of the Z-scored bulk RNA expression levels for some of the transcription factors (TFs) targeted in the screen in FACS-purified populations, using data from [23]. Hierarchical clustering, represented on the dendrogram, was performed across TFs. *PRDM16* and *SNAI2,* TFs also perturbed in our experiment, were not reported in this dataset. **(B)** Schematic of pooled single-cell screen genotyping for screen editing efficiency assessment using multiplexed single-cell PCR. **(C)** Box plots of the per cell percentages represented by each genotyping, colored by amplicon locus class. **(D)** Pseudo-bulk single-cell insertion, deletion, and substitution frequencies at one of the targeted RUNX1 loci, shown for control cells and cells with targeting sgRNAs. Mutational frequencies are a weighted frequency of the times that a given type of mutation was seen across reads for one cell barcode, which was then averaged across cells. Black dotted lines represent the boundaries of the sgRNA and the red dotted line represents the predicted cut site. **(E)** Scatter plot of the percentage of single-cells with at least one allele edited for each sgRNA from the pooled single-cell genotyping experiment and the average log2(average perturbation score) computed on RNA from the Perturb-multiome experiment. The Spearman correlation coefficient and a regression line with confidence intervals are shown. **(F)** Heatmaps of the scaled levels of the top differentially-accessible chromatin peaks between GATA1 sgRNA-targeting and control cells. Within each heatmap, each column represents a sample of single cells, ordered by the perturbation score. **(G)** Left: histograms of the flow cytometry measurements of intracellular TF SPI1 levels for HSPCs edited with AAVS1 control, SPI1_295, or SPI1_140 sgRNA. Right: histograms of the flow cytometry measurements of intracellular TF GATA2 pr for HSPCs edited with AAVS1 control, GATA2_562, or GATA2_176 sgRNA.

**Supplementary Figure 4.**
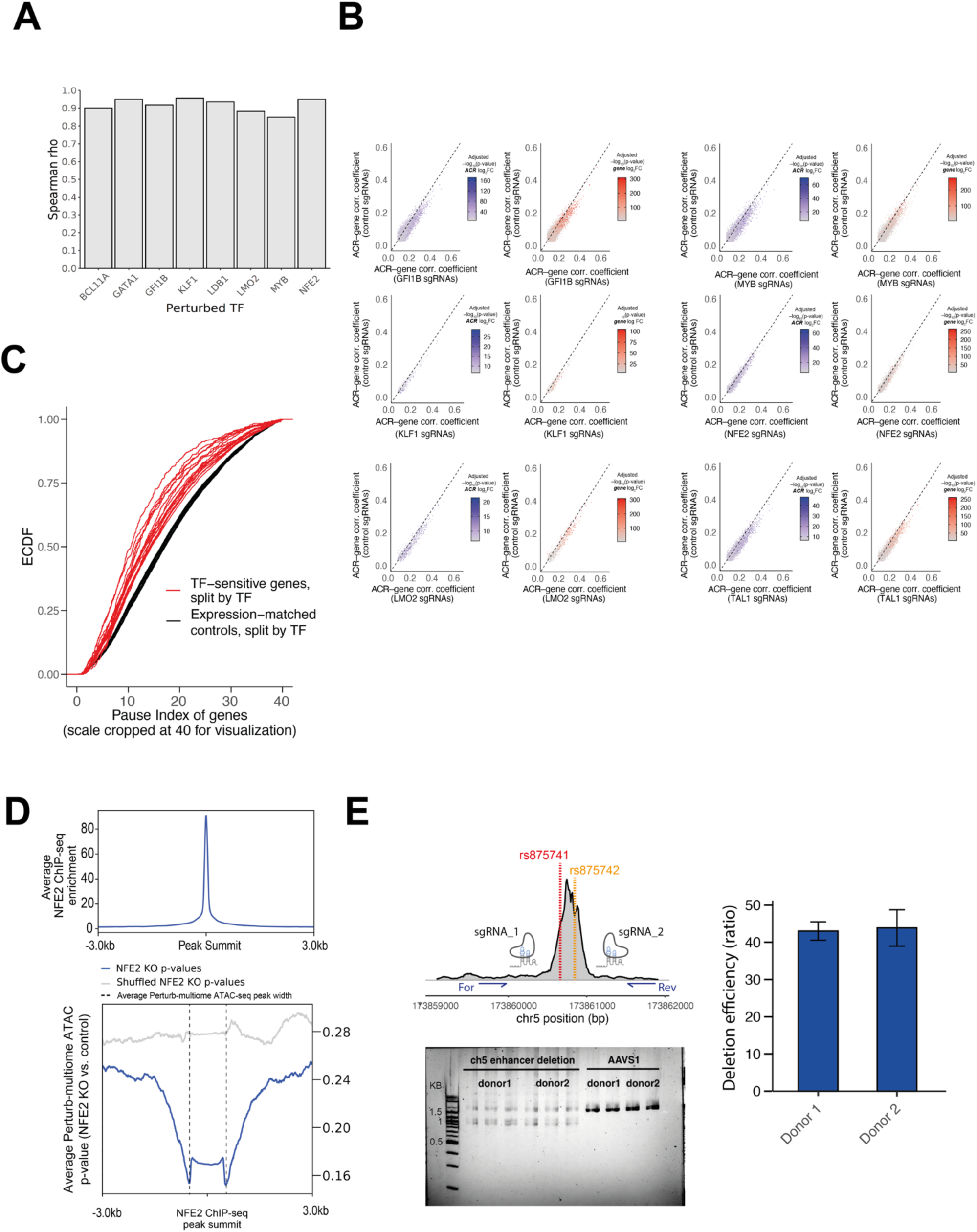
Properties of TF-sensitive genes and cis-regulatory elements and experimental validation of one of such linkages. **(A)** Spearman correlation coefficients of the gene expression model weights (i.e. the degree of perturbation of each gene) in two biological replicates that each used different HSPC donors and sgRNAs targeting different locations of the TF coding region. Data shown for TFs perturbed in both replicates. **(B)** Scatter plots of ACR–gene correlations across cells transduced with control sgRNAs and cells transduced with TF-targeting sgRNAs, for erythroid TF with more than 250 TF-sensitive ACR-gene pairs within 50kb from one another. The diagonal represents the correlation values for ACR–gene pairs with equal correlation coefficients among cells transduced with control sgRNAs and cells transduced with TF-targeting sgRNAs. Left panels: the color scale represents the p-value adjusted for multiple hypothesis testing of the ATAC ACR log2FC between TF-targeting and control sgRNAs. Right panels: the color scale represents the p-value adjusted for multiple hypothesis testing of the gene expression log2FC between TF-targeting and control sgRNAs. **(C)** Cumulative distribution plot of the pause index calculated with PRO-seq **(Materials and Methods)** of TF-sensitive genes for each TF (red) and control expression matched genes for each (black). **(D)** Top: profile plot of independent NFE2 ChIP-seq enrichment surrounding called ChIP-seq peaks from GYPA^+^ human erythroid precursors [56]. Bottom: profile plot of Perturb-multiome adjusted -log10(pvalues) for the log2FC of accessibility for peaks in cells with NFE2-targeting sgRNAs compared to control, centered on NFE2 ChIP-seq peaks as defined in the top panel. **(E)** Top left, schematic of the location of the two sgRNA targeting the GATA1-perturbation-sensitive, CPEB4 correlated ACR. The two vertical lines highlight two variants with pleotropic associations to blood cell traits. Bottom left, PCR products for primers flanking the CPEB4 ACR, shown for edited cells and AAVS1-edited controls, respectively. Data shown for two donors. Right, deletion efficiency ratio calculated from the area-under-the-curve using Agilent bioanalyzer on the PCR products from the CPEB4 enhancer deletion.

**Supplementary Figure 5.**
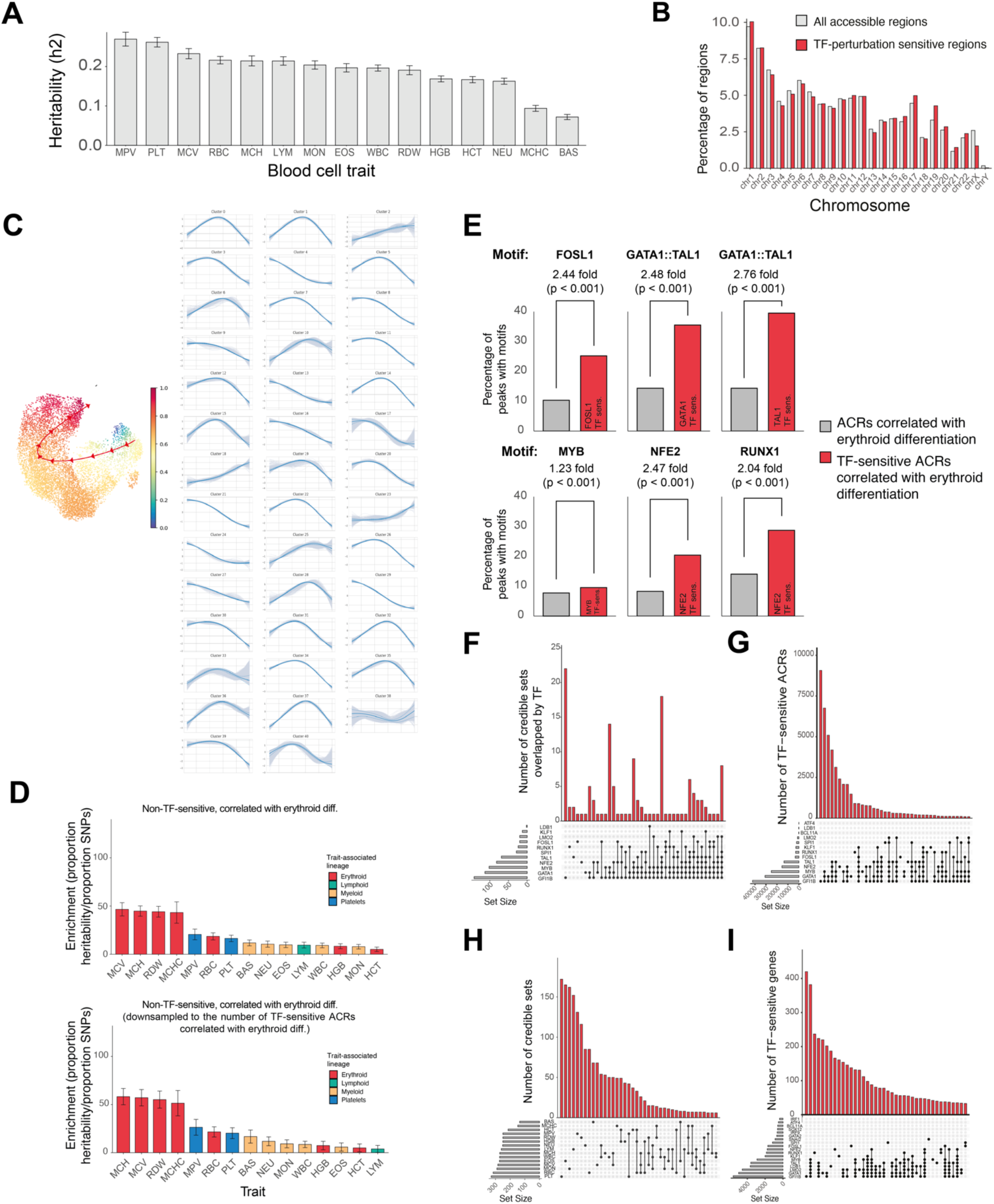
Characterization of trait-relevant variant enrichment in accessible chromatin regions with Perturb-multiome. **(A)** Percentage of heritability explained by genetic variation for blood cell traits **(Materials and Methods). (B)** Percentage of all accessible regions or TF-perturbation sensitive regions that is present in each chromosome. **(C)** Left, ATAC UMAP of Perturb-multiome used as input for Palantir to identify ACRs correlated with erythroid differentiation, colored by pseudotime **(Materials and Methods).** The red line shows the trajectory relative to which the dynamics of accessibility changes were identified. Right, clusters of ACR dynamic behaviors over the course of the aforementioned trajectory. Those that increased monotonically over the course of differentiation were defined as correlated with erythroid differentiation **(Materials and Methods). (D)** Top, barplots of the enrichment (proportion of heritability divided by the proportion of SNPs) for non-TF-sensitive ACRs correlated with erythroid differentiation, as defined in Fig. 4C. Bottom, barplots of the enrichment of non-TF-sensitive ACRs correlated with erythroid differentiation downsampled to the same number of TF-sensitive ACRs correlated with erythroid differentiation (a control for LDSC). Traits are colored by the respective associated lineage. **(E)** Barplots of the enrichment in TF-binding motifs between TF-sensitive ACRs correlated with erythroid differentiation and ACRs correlated with erythroid differentiation alone, regardless of their TF-sensitivity status. **(F)** Upset plot of the overlap between the credible sets overlapped by each transcription factor. **(G)** Upset plot of the overlap between the TF-sensitive ACRs arising from the perturbation of each transcription factor. **(H)** Upset plot of the overlap between the credible sets of each blood cell trait. **(I)** Upset plot of the overlap between the TF-sensitive genes arising from the perturbation of each transcription factor.

**Supplementary Figure 6.**
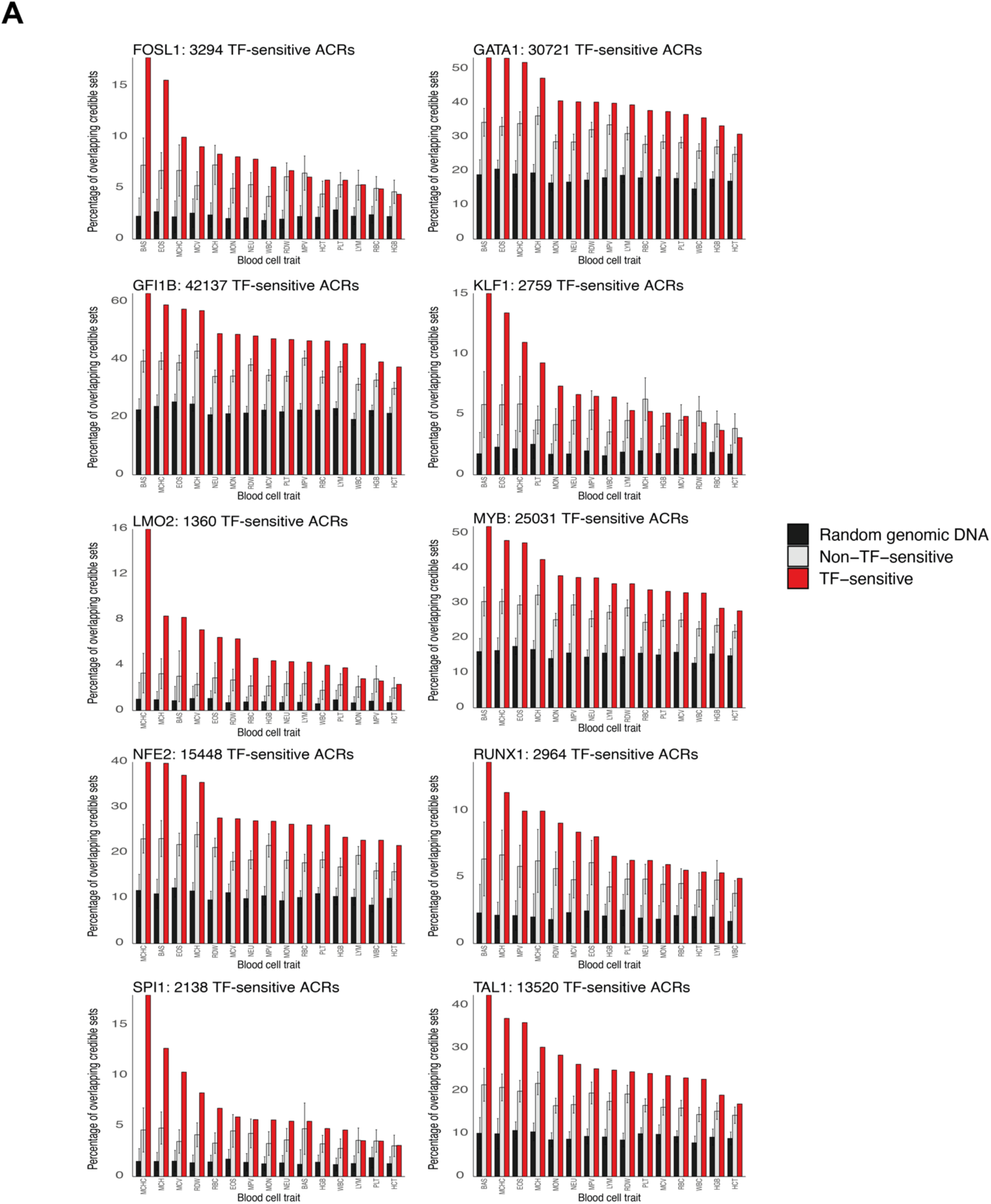
Overlap of blood-cell trait credible sets with TF-sensitive ACRs by TF. **(A)** Barplots for the percentage of 95% credible sets overlapping TF-sensitive ACRs, for each blood cell trait and TF, for TFs with more than 250 TF-sensitive ACRs. Error bars represent the standard deviation of 100 sampling events of non-TF-sensitive accessible ACRs (“non-TF-sensitive”) or of any genomic region (“random genomic DNA”) **(Materials and Methods).**

