## Supplementary Protocol for "Transcription factor networks disproportionately enrich for heritability of blood cell phenotypes"

**Step by step protocol for CROP-seq transcript PCR enrichment**

- Prep master mix for PCR1

PCR 1 reaction:

NEBNext Q5U master mix	15uL

F_CROPseq_PCR1, 25uM	1.25uL

AAO272, 25uM	1.25uL

Water	11.5uL

10X cDNA at 10ng/uL	1uL

Total volume	30uL

**Note:** starting with a 30-50ng/uL concentration might increase library complexity depending on the sample.

F_CROPseq_PCR1: /5Biosg/UAUAGTGACTGGAGTTCAGACGTGTGCTCTTCCGATC**TCGATTTCTTGGCTTTATATATCTTGTG**

AAO272: CTACACGACGCTCTTCCGAT*C*T

**Note:** KAPA HiFi Uracil+ Ready Mix works equally well - adjust annealing temperature accordingly.

**Note:** cDNA input amount might require optimization depending on the expression levels in the cell type.

PCR1

98C for 30s

Repeat 8 cycles:

i. 98C for 15s
ii. 69C for 15s
iii. 72C for 20s

72C for 120s

Hold at 4C

**Note:** Number of PCR1 cycles might require optimization depending on the expression levels in the cell type.

Perform SPRI purification using 1X beads by adding 30 uL of SPRIselect beads, thoroughly mixing with a pipette, and allowing it to incubate for a duration of 5 minutes. Afterwards, position it on position HIGH of the magnet and dispose of the supernatant. Follow this by washing twice using ethanol, centrifuge to concentrate, and eliminate any residual ethanol. Elute in 24µL of buffer EB. **The protocol can be stopped here at -20C.**

**Perform biotin pull-down using Dynabeads MyOne streptavidin beads as per invitrogen protocol, reproduced below:**

Prep **2X BW buffer**; see recipe from the manufacturer :

10 mM Tris-HCl (pH 7.5)

1 mM EDTA

2 M NaCl

1. Prepare Dynabeads MyOne streptavidin beads C1. Perform this cleanup in a single 1.5mL eppendorf for all your reactions (multiply the volumes below by your number of reactions).
   1. Prep **2X BW buffer** with the protocol above
   2. Add 12uL of Dynabeads.
   3. Add 50uL of **1X BW buffer**. Pipette mix.
   4. Place on magnet (high position) and discard supernatant
   5. Resuspend with 50uL 1x BW.
   6. Repeat steps 4 and 5 3 more times for a total of 4 washes
2. Add 24ul **2x BW buffer** to washed Dynabeads. Distribute into as many PCR strip wells as reactions that you have. Then, add to each the 24uL of PCR product.
3. Incubate for **15 mins** at **RT** on **hula mixer.**
4. Place on magnet (high), remove supernatant. Wash beads three times with 50µl **1X BW buffer** (note: pipette mix each wash to resuspend the beads). **Keep the beads.**
5. Resuspend in **15uL trisEDTA (TE)** + **0.5uL NEB USER enzyme**. Incubate for **30 minutes** (incubation can safely be reduced from 2 hours) at 37degC on ***hula mixer.***
6. Place on magnet (low), **transfer the supernatant** to a clean PCR tube.
7. Perform **1X SPRI cleanup** (add 15.5 uL of SPRIselect beads, pipette mix and incubate for 5 minutes, magnet and discard supernatant, wash 2x with 200µl 80% ethanol, spin down and remove any remaining ethanol). Elute in 15uL Elution Buffer (Qiagen) or H2O, incubate 2min at RT, place on magnet low and transfer the supernatant in another PCR strip.

Perform **PCR2, 27 cycles. This should be optimized for your cell type/experiment/MOI using qPCR: you can use 2.5uL of the prior elution for qPCR. We find 27-28 cycles is optimal for HSPCs.**

| NEB Q5 MASTER MIX (NEB, M0492L) | 25uL |
| --- | --- |
| Universal Illumina adapter P5, **100uM**  AATGATACGGCGACCACCGAGATCTACACTCTTTCCCTACACGACGCTC | 0.5uL |
| Universal Illumina adapter P7, **100uM** CAAGCAGAAGACGGCATACGAGATXXXXXXXXGTGACTGGAGTTCAGACGTGTGCTCTTCCGATCT  => Add a diff index for each reaction | 0.5uL |
| Clean PCR1 product | 12.5uL |
| Water | 11.5uL |
| Total reaction | 50uL |

**Enrichment_PCR2** : Run the following PCR:

98C for 30s

Repeat 27-28 cycles:

i. 98C for 15s
ii. 69C for 15s
iii. 72C for 20s

72C for 120s

Hold at 4C

Perform 1X SPRI cleanup (**50uL of beads**) and elute in 30uL of EB buffer.

Run 1µl of each PCR reaction on a gel to make sure it amplified

Pool 2µl of each PCR for the final library pool, then dilute 1:10 and Run BioA.

**Illumina Sequencing**

28 cycles R1 (cell barcode and UMI), 8 cycles I1 (sample index), 20 cycles R2 (sgRNA sequence, use Custom_sequencing_primer_R2_CROPseq

CGATTTCTTGGCTTTATATATCTTGTGGAAAGGACGAAACACCG
