## Supplementary Table S4 for "Transcription factor networks disproportionately enrich for heritability of blood cell phenotypes"

**Table S4. Multiparametric Flow Cytometry Antibodies**

| **Antibody** | **Manufacturer** | **Reference** | **Dilution** |
| --- | --- | --- | --- |
| PE anti-human CD34 | BioLegend | 343506 | 1:50 |
| FITC anti-human CD36 | BioLegend | 336204 | 1:50 |
| PE/Cy7 anti-human CD123 | BioLegend | 306010 | 1:50 |
| Brilliant Violet 421 anti-human CD71 | BioLegend | 334122 | 1:150 |
| APC anti-human CD235a | BioLegend | 349114 | 1:50 |
| 7-AAD Viable staining solution | BioLegend | 420404 | 1:50 |
| PE anti-human GATA2 | BioLegend | 614003 | 1:100 |
| PE anti-human SPI1 (PU.1) | BioLegend | 658009 | 1:100 |
